# Single-Cell Transcriptome Dynamics of the Autotaxin-Lysophosphatidic Acid Axis During Muscle Regeneration Reveal Proliferative Effects in Mesenchymal Fibro-Adipogenic Progenitors

**DOI:** 10.1101/2022.07.02.498539

**Authors:** Osvaldo Contreras, Richard P. Harvey

## Abstract

Lysophosphatidic acid (LPA) is a growth factor-like bioactive phospholipid recognising LPA receptors (LPARs) and mediating signalling pathways that regulate embryonic development, wound healing, carcinogenesis, and fibrosis, via effects on cell migration, proliferation and differentiation. Extracellular LPA is generated from lysophospholipids by the secreted hydrolase - ectonucleotide pyrophosphatase/phosphodiesterase 2 (ENPP2; also, AUTOTAXIN / ATX) and metabolised by different membrane-bound phospholipid phosphatases (PLPPs). Here, we use public bulk and singlecell RNA sequencing datasets to explore the gene expression of *Lpar*_1-6_, *Enpp2*, and *Plpp* genes under skeletal muscle homeostasis and regeneration conditions. We show that the skeletal muscle system dynamically expresses the *Enpp2-Lpar-Plpp* gene axis, with *Lpar1* being the highest expressed member among LPARs. *Lpar1* was expressed by mesenchymal fibro-adipogenic progenitors (FAPs) and tenocytes, whereas FAPs mainly expressed *Enpp2*. Clustering of FAPs identified populations representing distinct cell states with robust *Lpar1* and *Enpp2* transcriptome signatures in homeostatic cells expressing higher levels of markers *Dpp4* and *Hsd11b1*. However, tissue injury induced transient repression of *Lpar* genes and *Enpp2*. The role of LPA in modulating the fate and differentiation of tissue-resident FAPs has not yet been explored. Ex vivo, LPAR1/3 and ENPP2 inhibition significantly decreased the cell-cycle activity of FAPs and impaired fibro-adipogenic differentiation, implicating LPA signalling in the modulation of the proliferative and differentiative fate of FAPs. Together, our results demonstrate the importance of the ENPP2-LPAR-PLPP axis in different muscle cell types and FAP lineage populations in homeostasis and injury, paving the way for further research on the role of this signalling pathway in skeletal muscle homeostasis and regeneration, and that of other organs and tissues, in vivo.

**Summary:** Our reanalysis of single-cell transcriptomics revealed the involvement and temporally dynamic expression of the ENPP2-LPAR-PLPP axis in response to skeletal muscle regeneration.

## Introduction

Striated skeletal muscle is an endocrine organ regulating whole-body metabolism, heat, posture, and movement. This highly plastic tissue changes and adapts its function throughout an organism’s lifespan, making it an essential organ to maintain whole-body homeostasis. Mammalian adult skeletal muscle regeneration remains one of the most captivating and remarkable faculties in mammals (Baghdadi and Tajbakhsh, 2018). Although regenerative muscle capability relies on tissue-resident adult unipotent muscle stem cells (MuSCs, also known as satellite cells) (Lepper et al., 2011; Murphy et al., 2011; Sambasivan et al., 2011; Fry et al., 2015), recent discoveries have demonstrated that successful muscle regeneration requires a complex interplay between different cell types [reviewed in (Theret et al., 2021)]. Although significant progress has been made in understanding skeletal muscle regeneration, there is a need to identify novel, potentially druggable, targets to boost muscle repair in myopathies, neuromuscular disorders, muscle trauma and unhealthy aging. Fibro-adipogenic progenitors (FAPs) have recently emerged as essential stromal cells for maintaining skeletal muscle homeostasis, mass, neuromuscular integrity, and proper tissue regeneration [reviewed in (Giuliani et al., 2021; Theret et al., 2021)]. However, FAPs have also been proven to drive muscle degeneration, mediating exacerbated fibrous-adipose-bone ectopic deposition in severe trauma and myopathies [reviewed in (Contreras et al., 2021b; Molina et al., 2022)].

Lysophosphatidic acid (LPA, also known as lysophosphatidate) is a small circulating bioactive phospholipid [430-480 Da, equivalent to 4-5 amino acids] with a core that has a phosphate group, glycerol, and a fatty acid chain (Moolenaar, 1995; Okudaira et al., 2010). LPA can act as an extracellular signalling molecule via autocrine, paracrine, or endocrine processes (Moolenaar, 1995; Geraldo et al., 2021). Among its wide range of biological functions, LPA regulates platelet aggregation, smooth muscle cell contraction, cell differentiation, cell proliferation and survival, chemotaxis, carcinogenesis, and stem cell biology (van Corven et al., 1989, 1992; Fang et al., 2000; Binder et al., 2015; Lidgerwood et al., 2018; Magkrioti et al., 2018; Geraldo et al., 2021; Xu et al., 2021). LPA signalling is mediated by at least six different receptors (LPA_1-6_) encoded by individual genes, which recognise extracellular LPA species (Kihara et al., 2014; Geraldo et al., 2021). These receptors are members of the seven-transmembrane G protein-coupled receptors (GPCRs) superfamily, Class A rhodopsin-like and lipid-like receptors (Kihara et al., 2014). As such, LPARs signal through several intracellular effector pathways activated by heterotrimeric G proteins, including G_i/o_, G_12/13_, G_q/11_, and G_s_ [reviewed in (Geraldo et al., 2021)].

Extracellular levels of LPA are mainly controlled by the lysophospholipase D activity of the secreted enzyme ENPP2 (also known as AUTOTAXIN / ATX) (Akira et al., 1986; Tokumura et al., 2002; Okudaira et al., 2010). ENPP2 generates LPA by hydrolysis of lysophospholipids (lysophosphatidylcholine, lysophosphatidylserine and lysophosphatidylethanolamine), making it an essential enzyme for production and maintenance of extracellular and serum LPA (Umezu-Goto et al., 2002; Benesch et al., 2015). ENPP2 is required for proper mammalian development and *Enpp2*-null mice die around embryonic day 10 (Tanaka et al., 2006). Although ubiquitously expressed in adult tissues (Ninou et al., 2018), recent studies have shown ENPP2 expressions in adipose tissue as a major source of circulating and extracellular levels of this enzyme (Dusaulcy et al., 2011; Nishimura et al., 2014), suggesting that ENPP2 could act as an essential long and short distance adipokine (Funcke and Scherer, 2019).

Extracellular LPA is primarily metabolized by the ecto-activities of at least three plasma membrane-bound magnesium-independent lipid phosphate phosphatases or phospholipid phosphatase: PLPP1, PLPP2, and PLPP3, encoded by their respective *Plpp* genes [reviewed in (Brindley et al., 2009; Tang et al., 2015)]. However, other magnesium-independent phospholipid phosphatases with broader substrate specificity can also metabolize LPA, including PLPP4, PLPP5, and PLPP6 (Tang and Brindley, 2020). PLPP7 has no demonstrable enzymatic activity, and little information is available (Tang and Brindley, 2020). PLPPs catalyze the dephosphorylation of various glycerolipid and sphingolipid phosphate esters, regulating their bioavailability (Brindley et al., 2009). Because of their crucial role in metabolizing LPA, gaining knowledge about the gene expression dynamics and regulation of ENPP2 and PLPPs, and their associated genes, could bring novel interventional strategies for treating disease.

Accumulative evidence suggests the participation of the ENPP2-LPA-LPAR axis in skeletal muscles. Yoshida and colleagues provided the first evidence demonstrating that LPA acts in skeletal muscle cells. These authors showed *in vitro* that LPA induced C2C12 myoblast proliferation and cell growth while inhibiting myotube differentiation through Gi proteins (Yoshida et al., 1996). Interestingly, structurally related lipids did not exert the same pro-proliferative and anti-fusion effects as LPA or phosphatidic acid (PA) (Yoshida et al., 1996). Initial evidence suggested the expression of some *Lpar* genes in C2C12 myogenic cells in which LPA activates two known pro-mitogenic signalling pathways, ERK1/2 and AKT (Jean-Baptiste et al., 2005). Other supporting studies have shown that LPA increases intracellular Ca^2+^ concentration and induces DNA synthesis (Xu et al., 2008), reinforcing that LPA modulates myogenic cell growth and proliferation (Bernacchioni et al., 2018). Recently, Gomez-Larrauri and colleagues reported that PA induces DNA synthesis in C2C12 myoblast via LPAR1/LPAR2 and downstream ERK1/2-AKT signalling at similar concentrations to LPA (Gomez-Larrauri et al., 2021). Pharmacological inhibition of ENPP2 or *Enpp2* knockdown reduces myogenic differentiation, via a mechanism whereby *Enpp2* is a direct target gene of WNT/RSPO2-mediated TCF/LEF/β-CATENIN signalling (Sah et al., 2020). The authors also showed that whole-body conditional deletion of *Enpp2* impairs muscle regeneration upon acute BaCl_2_-induced damage (Sah et al., 2020). Reasoning in favour of a myogenic and pro-regenerative role for ENPP2, Ray and colleagues recently revealed that the ENPP2 axis regulates skeletal muscle regeneration in a satellite cell-specific manner (Ray et al., 2021). Thus, cumulative evidence shows that the ENPP2-LPAR axis is active in striated muscles modulating muscle stem cell function, adult myogenesis, hypertrophic muscle growth, and regeneration.

Because the exploration of the ENPP2-LPAR-PLPP network in muscles has been highly limited to myogenic and satellite cells, there is a current lack of knowledge about the gene expression dynamics of this axis in other muscle cells in response to injury. Here, utilizing publicly available bulk RNA-seq and single-cell transcriptomic (scRNA-seq) datasets, we studied for the first time the gene expression and pathway dynamics of the ENPP2-LPAR-PLPP network and its changes in numerous cell types in adult muscle homeostasis and regeneration, including subsets of immune cells, muscle stem cells, tenocytes, and fibro-adipogenic progenitors. In addition, we compared the effects of two specific pharmacological inhibitors of LPAR1/3 (Ki16425) and ENPP2 (PF-8380) in modulating cell growth, proliferation, and fibro-adipogenic differentiative fate on adult mesenchymal FAPs and satellite cells.

## Results

### Skeletal muscle differentially expresses ENPP2-LPAR-PLPP coding genes

To study ENPP2-LPAR-PLPP pathway gene expression dynamics in adult skeletal muscle tissue, we utilized public bulk transcriptomic data (Scott et al., 2019) and evaluated ENPP2-LPAR-PLPP gene expression in different samples: whole muscle, lineage^+^ cells (CD31^+^/CD45^+^), lineage^-^ cells (CD31^-^/CD45^-^), and Lineage^-^/SCA1^+^ FAPs (**Fig. 1A**). In whole muscle tissue, genome-wide transcriptomic profiling showed differential expression of LPAR members. *Lpar1* was the most expressed member, followed by *Lpar6* and *Lpar4*, whereas *Lpar2, Lpar3*, and *Lpar5* levels were very low (**Fig. 1B**). Limb muscle also expresses *Enpp2* (~8 FPKM or fragments per kilobase of exon per million mapped fragments) (**Fig. 1B**). Moreover, we evaluated *Plpp* gene expression dynamics in skeletal muscle tissue. *Plpp1, Plpp3*, and *Plpp7* were highly expressed compared to *Plpp2, Plpp4, Plpp5*, and *Plpp6*. Interestingly, *Lpar6*, *Plpp2*, and *Plpp6* were highly enriched in the lineage^+^ fraction, suggesting they may be expressed by endothelial or hematopoietic lineage (**Fig. 1A**). Thus, most ENPP2-LPAR-PLPP pathway components are present in healthy adult skeletal muscle.

**Figure 1.**
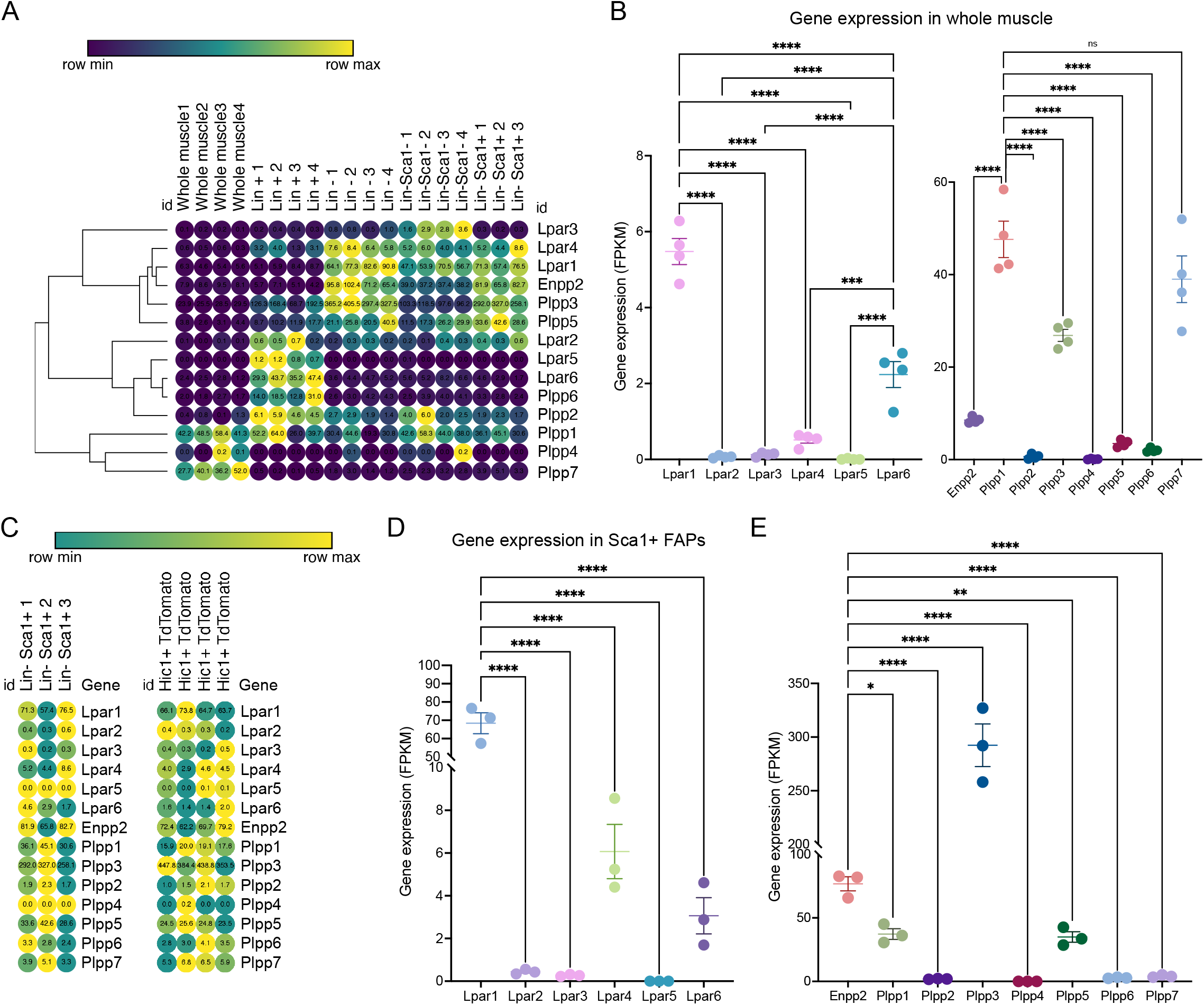
Bulk RNAseq transcriptomics analysis revealed differential gene expression of the LPA-LPAR-Autotaxin-Plpp network. (A) Heat map showing gene expression levels of *Lpar, Enpp2*, and *Plpp* genes in whole skeletal muscle tissue, Lineage^+^, Lineage^-^, and Lineage^-^Sca1^+^ FAPs from a bulk RNAseq library [Scott et al., 2019]. Gene expression is shown as fragments per kilobase of exon per million mapped fragments (FPKM). (B) Quantification of *Lpar, Enpp2*, and *Plpp* genes transcript abundance (FPKM) in muscle tissue. (C) Heat map showing gene expression levels of *Lpar, Enpp2*, and *Plpp* genes in Lineage^-^Sca1^+^ FAPs and Hic1 + tdTomato expressing cells [Scott et al., 2019]. (D) Quantification of Lpar_(1-6)_ genes transcript abundance (FPKM) in Sca1^+^ FAPs. (E) Quantification of *Enpp2* and *Plpp* genes transcript abundance (FPKM) in Sca1^+^ FAPs.

### SCA1^+^ fibro-adipogenic progenitors abundantly express ENPP2-LPAR-PLPP pathway genes in resting state

Since FAPs have a crucial role in regulating muscle and neuromuscular tissue integrity, we evaluated gene expression of the ENPP2-LPAR-PLPP gene network in uninjured muscle-resident SCA1^+^ FAPs and Hic1-lineage^+^ (tdTomato^+^) mesenchymal stromal cells (Scott et al., 2019). As observed in skeletal muscle tissue, LPA receptors were differentially expressed in resting FAPs (**Fig. 1C, D**). *Lpar1* was the most expressed family member, followed by *Lpar4* and *Lpar6*, respectively (**Fig. 1C, D**). However, *Lpar2, Lpar3*, and *Lpar5* were not significantly expressed in FAPs (**Fig. 1C, D**). These results indicate that *Lpar1* is the highest expressed LPAR member in stromal FAPs, as seen in fibroblast lineages in other tissues (**Fig. S1**).

FAPs express relatively high levels of *Enpp2* (~77 FPKM) (**Fig. 1E**), suggesting FAPs could be a significant cell source of extracellular LPA. Of the *Plpp* genes, *Plpp3* was the highest expressed member, followed by *Plpp1* and *Plpp5. Plpp2, Plpp4, Plpp6*, and *Plpp7* genes were very low expressed (**Fig. 1E**). The trend of ENPP2-LPAR-PLPP pathway gene expression is similar between SCA1^+^ FAPs and *Hic1-lineage^+^* FAP cells (**Fig. S2**). These results show that ENPP2-LPAR-PLPP pathway genes are significantly expressed in FAPs and, therefore, suggest a role for the bioactive phospholipid LPA and LPA-mediated signalling in skeletal muscle and stromal progenitor cells in homeostasis.

### Analysis of ENPP2-LPAR-PLPP network gene expression in skeletal muscle using single-cell transcriptomics

To gain more detailed insights into the role of the LPA axis in adult skeletal muscle cells, we further evaluated the relative expression of its network genes in skeletal muscle cells using scRNA-seq data (Oprescu et al., 2020). The single-cell data was derived from uninjured and injured muscle sampled at different time points from early hours post-injury to damage resolution (**Fig. 2A; Fig. S3**). Here, we identified cells with discrete lineage identities and transcriptional states, performing unbiased clustering on an aggregate of cells using the *Seurat* R package (Butler et al., 2018) (**Fig. 2A; Fig. S3**). We initially obtained 29 distinct clusters across different conditions (**Fig. S3A-C**). We visualise distinct cell populations in UMAP dimensionality reduction plots (Materials and Methods), representing a total of 19 cell populations and 7 distinct cell lineages across uninjured and injured conditions (**Fig. 2A; Fig. S4**).

**Figure 2.**
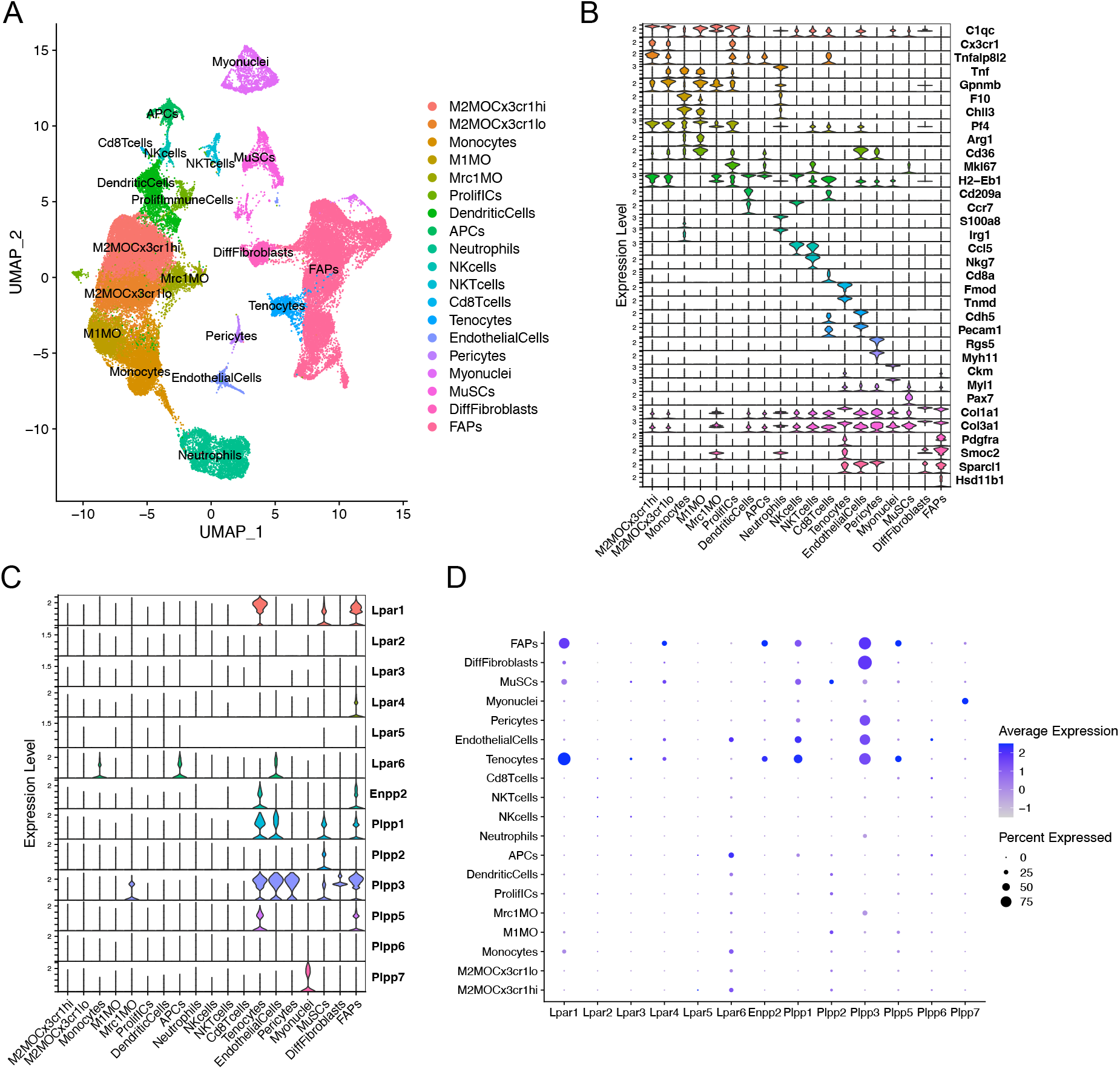
Analysis of LPAR-Autotaxin/Enpp2-PIpp gene expression at single-cell resolution. (A) Uniform manifold approximation and projection (UMAP) plot of scRNA-seq public data [Oprescu et al., 2020] showing 19 distinct cell lineages in single cells across skeletal muscle homeostasis and regeneration. Detected major cell lineages were colored by the predominant cell type(s) that composes each cluster. (B) Violin plots showing the expression level of several marker genes across the different populations depicted in (A). (C) Violin plots showing the gene expression level of LPAR, Enpp2, and Plpp family members across the different populations or cell clusters. (D) Dot plot showing gene expression levels of LPAR, Enpp2, and Plpp family members. Dot plots help to visualize two values across two dimensions: color and size. The color gradient of the dot approximates average gene expression (light grey: low expression; Navy blue: high expression). Dot size represents the

Major cell types and their defining marker signatures comprised fibro-adipogenic progenitors (FAPs; *Pdgfra^+^Pi16^+^Smoc2^+^Hsd11b1^+^Cxcl14^+^*), differentiated fibroblasts (DiffFibroblasts; *Pdgfra^-^Meg3^+^Lum^+^Col1a1^+^Dlk1^+^*), muscle stem cells/satellite cells (MuSCs; *Cdh15^+^Pax7^+^Myog^+^Megf10^+^*), myonuclei (*Ttn^+^Ckm^+^Myh1^+^*), pericytes (*Rgs5^+^*, which also includes markers of smooth muscle cells i.e., *Myh11*), endothelial cells (*Pecam1^+^Cdh5^+^Kdr^+^Cd36^+^*), tenocytes (*Tnmd^+^Mkx^+^Fmod^+^Kera^+^*), CD8^+^ T cells (*Cd8a^+^*), natural killer T cells (NKTcells; *Nkg7^+^*), natural killer cells (NKcells; *Ccr7^+^Ccl5^+^*) neutrophils (*S100a8^+^S100a9^+^Irg1^+^Tnf^+^*), antigen presenting cells (i.e., B cells, among others) (APCs; *H2-Eb1^+^*), dendritic cells (DCs; *Cd209a^+^H2-Eb1^+^Ccr7^+^*), proliferative immune cells (ProlifICs; *Stmn1^+^Birc5^+^Mki67+Acp5^+^*), Mrc1 macrophages (Mrc1MO; *Mrc1^+^C1qc^+^Cx3cr1^-^Gpnmb^-^*), M1 macrophages (M1MO;*Cx3cr1^-^Pf4^+^Arg1^+^Cd36^+^*), monocytes (*F10^+^Chil3^+^Tnf+*), and two related M2 macrophages (M2MO) - M2MOCx3cr1hi (*C1qc+Cx3cr1^hi^Tnfaip8l2^hi^Gpnmb^low^*) and M2MOCx3cr1lo (*C1qc^+^Cx3cr1^low^Tnfaip8l2^low^Gpnmb^hi^*) (**Fig. 2A, B; Fig. S4**).

Within the FAP lineage, we observed high transcriptional variation and identified several cluster subtypes (**Fig. S3A-C; Fig. S4**). However, for preliminary analyses involving major cell lineages we intentionally grouped the 7 main FAP subclusters (clusters 12, 8, 2, 9, 4, 20, and 21) and kept differentiated fibroblasts (DiffFibroblasts, cluster 15) aside (**Fig. 2A, B; Fig. S3; Fig. S4**). DiffFibroblasts have a differentiated fibroblasts/myofibroblast-like gene signature, highly expressing markers of activation and differentiation, and loss of stemness markers (*Pdgfra^-^Pi16^-^Lum^+^Col1a1^+^Dlk1^+^Col3a1^+^Col6a3^+^*) (**Fig. 2B; Fig. S4**). Specifically, downregulation of *Pdgfra* has been shown to be a sign of a differentiated FAP phenotype and correlates with their loss of stemness (Contreras et al., 2019a, 2020; Soliman et al., 2020).

Our analysis shows that fibro-adipogenic progenitors, tenocytes and MuSC/satellite cells predominantly express *Lpar1*, but its expression was higher in tenocytes and FAPs than MuSCs (**Fig. 2C, D**). *Lpar1* expression has not previously been shown in tenocytes or FAPs, although it has been reported that myogenic MuSCs express functional LPAR1 (Ray et al., 2021). *Lpar4* was expressed in FAPs but not highly expressed in other cell types (**Fig. 2C, D**). *Lpar2, Lpar3*, and *Lpar5* genes were virtually absent in FAPs (**Fig. 2C, D**), which corroborates our previous results exploring bulk RNAseq data of SCA1^+^ FAPs (**Fig. 1D**). On the contrary, *Lpar6* had a broader cell type-dependent expression, including in different populations of immune cells (e.g., monocytes and APCs) and endothelial cells (**Fig. 2C, D**). Interestingly, *Lpar6* is the only LPAR gene member expressed in the immune cell lineage, suggesting that LPA or related phospholipids may also modulate immune cell function. These findings better define the bulk RNAseq analyses shown in **Figure 1** for lineage^+^ cells.

FAPs and tenocytes expressed high levels of *Enpp2*, which was barely detected in other cell types (**Fig. 2C, D**). This suggests FAPs and tenocytes as the two major cell types responsible for extracellular LPA production in skeletal muscles. With relation to LPA catabolizing enzymes, FAPs highly expressed *Plpp1*, followed by *Plpp3* and *Plpp5* (**Fig. 2C, D**). MuSCs expressed *Plpp1* and *Plpp2*, but no other members, whereas pericytes only *Plpp3* (**Fig. 2C, D**). Endothelial cells highly expressed *Plpp3* and, to a lesser degree, *Plpp1* (**Fig. 2C, D**). Tenocytes also highly expressed *Plpp3* and *Plpp1*, and less *Plpp5* (**Fig. 2C, D**). M2-like MCR1^+^ macrophages specifically expressed *Plpp3*. Myonuclei only expressed *Plpp7* (**Fig. 2C, D**). Intriguingly, we could not detect *Plpp4* in the analyzed data, which could be due to the very low expression of this *Plpp* gene as seen exploring bulk RNAseq data (**Fig. 1**). Hence, our analysis reveals for the first time the detailed landscape of *Enpp2-Lpar-Plpp* gene expression in several muscle cell types in homeostasis and regeneration at single-cell resolution, suggesting an active role of FAPs and tenocytes in producing LPA.

### Analysis of ENPP2-LPAR-PLPP axis in fibro-adipogenic progenitor subpopulations in response to skeletal muscle regeneration

Fibro-adipogenic progenitors and their descendant lineages are the primary cell types responsible for ectopic fibrosis, fatty tissue, and bone formation and deposition in severe myopathies, degenerative disorders and neuromuscular disease (Contreras et al., 2021). Thus, we explored the temporal gene expression dynamics of *Enpp2-Lpar-Plpp* gene members in FAPs in homeostasis and in response to acute injury. We performed unbiased clustering on an aggregate of the initial clusters 12, 8, 2, 9, 4, 20, and 21 to increase the resolution of our fibro-adipogenic progenitor analyses (**Fig. S3**). We decided to include tenocytes (initial cluster 17 (**Fig. S3; Fig. S4**)) in our clustering analysis since these cells share a mesenchymal origin and highly express *Lpar1* and *Enpp2*.

Our unbiased subcluster analysis retrieved 10 distinct clusters (**Fig. 3A**). Using *FindAllMarkers* Seurat function on these clusters and determining the top 8 marker genes, we assigned different names for the eight FAP subtypes obtained and tenocytes (**Fig. 3B, C**). All FAP subpopulations showed expression of canonical FAP markers *Pdgfra* and *Sparcl1*, albeit at varying proportions and levels (**Fig. S5A-C**), and major changes in cell proportions were seen between conditions or days of injury (**Fig. 3D, E**). We named these cells Sparcl1 FAPs, Csrp2 FAPs, Dlk1 FAPs, Dpp4 FAPs, Hsd11b1 FAPs, Tyrobp FAPs (previously named as DiffFibroblasts), Cycling FAPs and Ccl2 FAPs, starting from the most numerous subpopulations to the less abundant (**Fig. 3B, C**). We also observed noticeable transcriptomic changes in the top 8 expressed genes following muscle injury (**Fig. 3D**). The top 8 expressing genes of Sparcl1 FAPs were *Sparcl1, Abca8a, Col15a1, Hmcn2, Htra3, Ltbp4, Penk* and *Cfh* (**Fig. 3E**). Csrp2 FAPs highly expressed *Csrp2, Sfrp2, Ltbp2, Lrrc15, Tnc, 1500015O10Rik, Acta2* and Tagln, whereas Dlk1 FAPs highly expressed *Dlk1, Igf2, Plagl1, Mest, Zim1, H19, Nrk* and *Agtr2*. The Dpp4 FAPs top 8 expressed genes were *Efhd1, Pcolce2, Dpp4, Sema3c, Cd55, Pi16, Efemp1* and *Stmn4. Hsd11b1, Ccl11, Crispld2, Vwa1, Enpp2, G0s2, Nmb* and *Inmt* genes characterized Hsd11b1 FAPs from other FAP subtypes, although these also highly express *Cxcl14* (**Fig. 3E**). Tyrobp FAPs expressed *Tyrobp, Fcer1g, Ctss, Lyz2, Laptm5, Slfn2, Cd52* and *Srgn*, whereas Cycling FAPs were characterized by high expression of genes related to survival and cell cycle, including *2810417H14Rik, Stmn1, Birc5, Mki67, Cks2, Tpx2, Cenpa* and *Top2a*. Finally, Ccl2 FAPs high expressed *Cxcl5, Ddx21, Ccl2, Rdh10, Slco2a1, Prg4, Lif* and *Mmp3*, highlighting a pro-inflammatory state of these cells at 12 hours post-injury (**Fig. 3E**). The tenocyte cluster highly expressed *Tnmd*, *Fmod*, *Thbs4*, *Col11a1*, *Cilp2*, *Scx*, *Kera* and *Chodl*, as previously described (Harvey et al., 2019; Scott et al., 2019).

**Figure 3.**
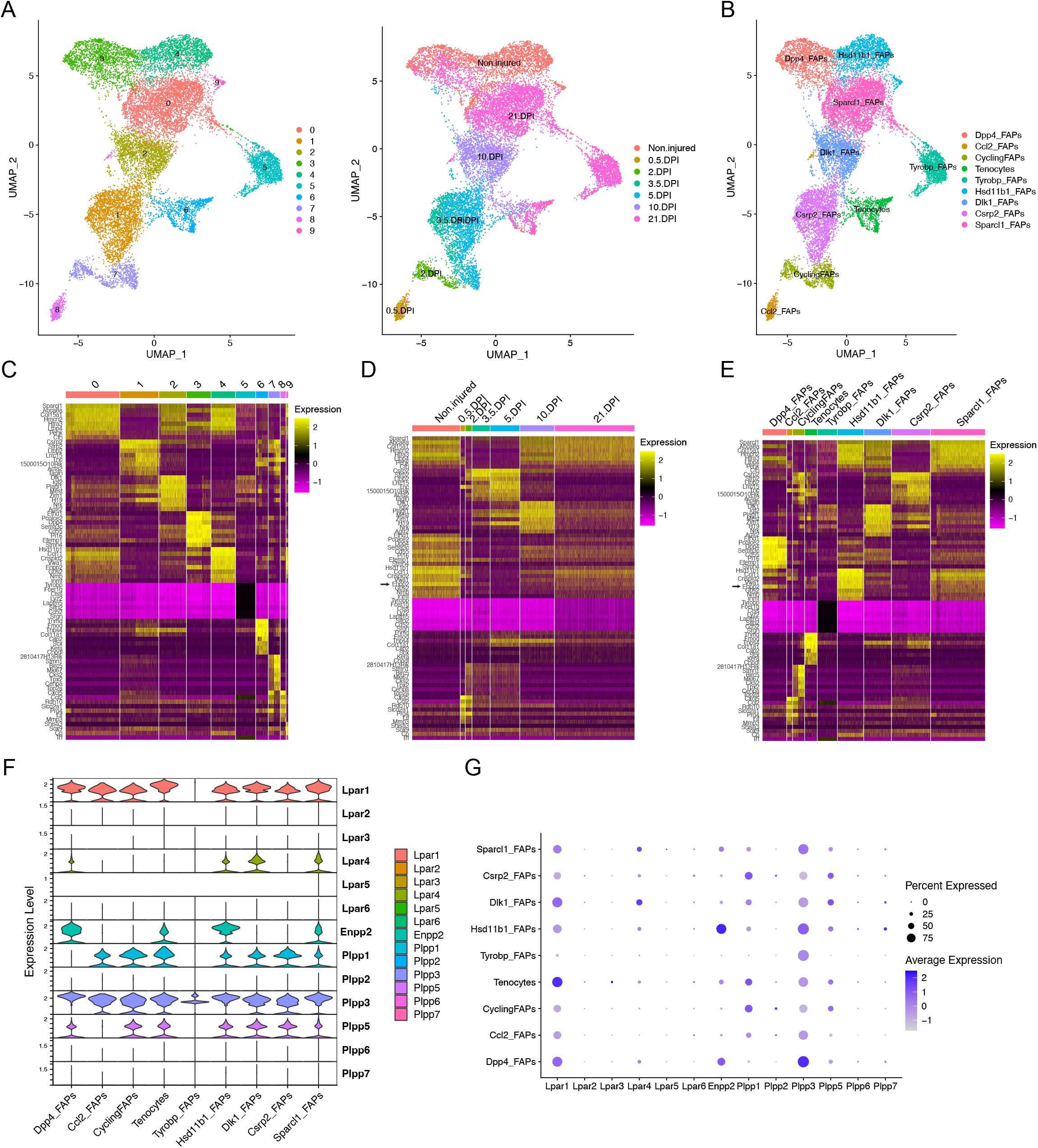
Resting fibro-adipogenic progenitors predominantly express LPA receptors and LPA-producing enzyme Enpp2/Autotaxin. (A) UMAP plot showing 10 distinct clusters across skeletal muscle homeostasis and regeneration when subclustering FAPs, DiffFibroblasts, and tenocytes subclusters. Detected major cell lineages and states were colored by the predominant cell type(s) that composes each cluster (0-9). (B) UMAP plot showing 9 distinct clusters (8 clusters for FAP lineage and 1 for tenocytes), which are named based on the most highly expressed gene shown in the heat maps shown below (C, E). (C) Heat map plot showing *top 8* expressed genes in each individual initial cluster shown in (A). (D) Heat map plot showing *top 8* expressed genes in the grouped 9 distinct clusters at different conditions (undamaged and days post-injury [DPI]). (E) Heat map plot showing the named clusters as described in the text, having its name because of one of the *top 8* expressed genes in each subset. (E) UMAP plot showing individual cells grouped based on the different conditions (uninjured and injured muscle) at different time points. DPI: days post-injury. (F) Violin plots showing the gene expression level of LPAR, Enpp2, and Plpp family members across the different FAPs and tenocytes subclusters. (G) Dot plot showing gene expression levels of LPAR, Enpp2, and Plpp family members. Note that Dpp4 and Hsd11b1 FAPs highly express *Enpp2/ATX* gene.

In uninjured conditions, we could distinguish two distinct FAP populations based on scRNA-seq, named Hsd11b1 FAPs, and Dpp4 FAPs after their highest upregulated genes (**Fig. 3A-E; Fig. S5**), as previously described (Scott et al., 2019; Oprescu et al., 2020). Early in the injury process, Ccl2 FAPs appear and relate to an activated immune-like pro-inflammatory FAP subpopulation mostly present at 12 hours post-injury (**Fig. 3A-B**). Cycling FAPs uniquely expressed a potent cell cycle gene signature, representing the most abundant FAP subtype found at 2 days post-injury (**Fig. 3A-E; Fig. S5**). Cycling FAPs can also be found at 3.5- and 5-days post-injury but to a lesser extent (**Fig. 3A-E; Fig. S5**). Csrp2 FAPs are more abundant at 3.5- and 5-days post-injury, whereas Dlk1 FAPs were present at 10 days (**Fig. 3A-E; Fig. S5**). The final captured stage of skeletal muscle regeneration, corresponding to day 21, mostly identified Sparcl1 FAPs together with Tyrobp FAPs (corresponding to DiffFibroblasts in our initial clustering) and, to a lesser extent, Dpp4 FAPs (**Fig. 3A-E; Fig. S5**).

Next, we further identified the expression profiles of the ENPP2-LPAR-PLPP axis in the different FAP subpopulations. Most major FAP subtypes expressed *Lpar1*, including Dpp4, Dlk1 and Sparcl1 FAPs at high levels (**Fig. 3F, G**). *Lpar1* was also expressed in Hsd11b1, Csrp2, Ccl2 and Cycling FAPs, although to a lesser extent (**Fig. 3F, G**). *Lpar1* was also highly expressed by tenocytes (**Fig. 3F, G**). Noticeable, *Lpar1* gene expression was undetectable in Tyrobp FAPs compared to the other 7 FAPs subtypes, suggesting LPAR1-dependent signalling may be downregulated in day 21 differentiated fibroblasts-like FAPs (**Fig. 3F, G**). No significant gene expression was detected for *Lpar2, Lpar3, Lpar5*, and *Lpar6* in FAPs or tenocytes (**Fig. 3F, G**). *Lpar4* was primarily expressed in Dlk1 and Sparcl1 FAPs and to a less extent in uninjured Hsd11b1 FAPs and Dpp4 FAPs (**Fig. 3F, G**). These results show that different FAP subpopulations that exist in homeostasis and those that appear following acute damage express different levels of LPA receptors, suggesting that LPA modulates FAP activation, survival, and fate primarily throughout LPAR1 and LPAR4. Also, the absence of LPA receptor gene expression in the Tyrobp FAPs subtype compared to their counterparts at day 21 (e.g., Sparcl1 and Dpp4 FAPs) suggests Tyrobp FAPs may be refractory to LPA actions (**Fig. 3E-G**). Thus, our single-cell exploration reports highly dynamic gene expression of LPA receptors in fibro-adipogenic progenitors in homeostasis and skeletal muscle regeneration.

*Enpp2* gene expression was higher in uninjured Hsd11b1 FAPs and Dpp4 FAPs than Sparcl1 FAPs, however, repressed in other FAP subpopulations (**Fig. 3E-G**). Remarkably, *Enpp2* expression was among the top 5 markers expressed in Hsd11b1 FAPs (**Fig. 3C, E**). Dpp4 FAPs also highly expressed *Enpp2* (**Fig. 3C-G**), suggesting a role for the encoded LPA extracellular-producing enzyme in resting FAPs, yet to be discovered. Tenocytes expressed *Enpp2* at levels comparable to Sparcl1 FAPs (**Fig. 3E-G**). Among *Plpp* genes, *Plpp3* gene expression was the most broadly distributed (Fig. 3E-G), although its expression was higher in Dpp4 FAPs, followed by Hsd11b1 and Sparcl1 FAPs, but was downregulated in other FAP subpopulations (**Fig. 3E-G**). *Plpp1* and *Plpp5* gene expression patterns were similar, except for Ccl2 FAPs that did not express *Plpp5*, only *Plpp3* (**Fig. 3E-G**). Among *Plpp* genes, Tyrobp FAPs only expressed one family member, *Plpp3* (**Fig. 3E-G**). As previously suggested, *Plpp2*, *Plpp6*, and *Plpp7* genes were absent in most FAP subtypes, with a small percent of Ccl2 FAPs expressing *Plpp2* and Hsd11b1 FAPs expressing *Plpp7* (**Fig. 3E-G**). Thus, our single-cell transcriptomics analysis showed enrichment for transcripts encoding the extracellular LPA-producing enzyme ENPP2 in two FAP subpopulations, Hsd11b1 and Dpp4, suggesting that resting FAPs as a significant source of LPA in skeletal muscles.

### Downregulation of LPA receptors in fibro-adipogenic progenitors in response to acute injury

To better understand the single-cell gene expression patterns of the LPA receptor family in adult FAPs during muscle regeneration, we grouped all FAP clusters. We then evaluated *Lpar* gene expression in response to injury (**Fig. S5E**). It was evident that skeletal muscle injury triggers a rapid but transient downregulation of LPA receptors in FAPs, including *Lpar1, Lpar4*, and *Lpar6* (**Fig. 4A; Fig. S5E**). *Lpar1, Lpar4*, and *Lpar6* gene expression was most noticeably downregulated at 12 h and stayed low up to 48 h post-muscle injury (**Fig. 4A; Fig. S5E**). Then, their expression increased towards pre-injury levels from 3.5 days to 10 days post-injury (**Fig. 4A**). Since changes happen when FAPs are activated and commit to proliferation and expand their numbers (**Fig. 3; Fig. S5D**), these data suggest an association between FAP activation and cell cycle dynamics during the period of repression of LPAR family genes. Finally, we determined the relative expression of LPA receptors genes at the genome-wide transcript level in quiescent and injury-activated *Hic1*-lineage^+^ (tdTomato^+^) mesenchymal stromal cells in muscle, found to be enriched in FAPs (Scott et al., 2019) (**Fig. 4B, C**). *Lpar1* was highly and preferentially expressed over *Lpar4* and *Lpar6* in quiescent and injury-activated *Hic1*-lineage^+^ cells (**Fig. 4B, C**). *Lpar2, Lpar3*, and *Lpar5* were more lowly expressed (**Fig. 4B, C**). *Lpar1* gene expression was early repressed in *Hic1*-lineage^+^ cells following acute damage, reaching its lowest on day 3 but recovering from day 4 post-injury onwards. Hence, two independent datasets demonstrate that the expression of LPAR gene family members is dynamically downregulated in injury-activated FAPs and *Hic1*-lineage^+^ cells but recovers later as muscle damage resolves through regeneration.

**Figure 4.**
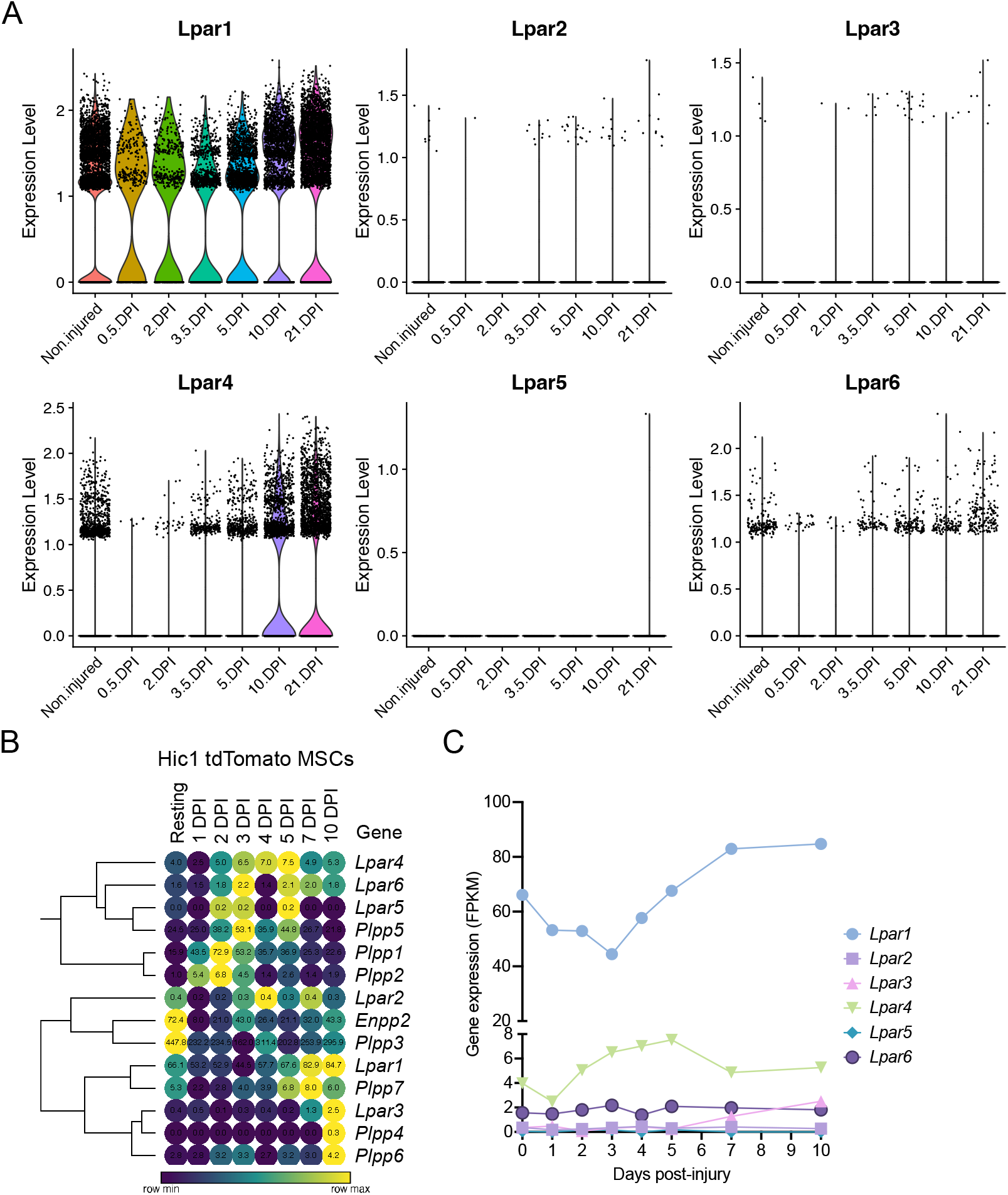
Muscle injury triggers a fast and strong downregulation LPA receptors in fibro-adipogenic progenitors. (A) Violin plots showing the gene expression level of LPAR family members and dynamics in response to injury. DPI: Days post-injury. (B) Heat map showing gene expression levels of Lpar, Enpp2, and Plpp genes in Hic1+ tdTomato expressing cells upon acute muscle damage [Scott et al., 2019]. (C) Quantification of Lpar_(1-6)_ genes transcript abundance (FPKM) in Hic1+ tdTomato expressing cells and dynamics in response to injury. DPI: Days post-injury.

### Repression of *Enpp2* and *Plpp* genes in fibro-adipogenic progenitors in response to acute injury

Next, we evaluated *Enpp2* expression in *Hic1*-lineage^+^ cells. *Enpp2* was highly expressed in quiescent *Hic1*-lineage^+^ cells but then sharply downregulated 1-day post-injury, before increasing again up to day 3, then reducing again until day 5 post-acute muscle injury (**Fig. 5A; Fig. S5E**). Expression increased again from day 5 to day 10 post-injury (**Fig. 5A**). At single-cell resolution, *Enpp2* showed an expression pattern in FAPs similar to that in *Hic1*-lineage^+^ cells (**Fig. 5A, B**), which is expected since *Hic1*-expressing cells mainly comprise FAPs in adult skeletal muscles (Scott et al., 2019; Contreras, 2020). Hence, *Enpp2* is downregulated to almost undetectable levels in FAPs at regenerative time points that associate with cell activation and proliferation (**Fig. S5D, E**). Levels remained very low up to 5 days, then recovered from day 10 to day 21 post-injury (**Fig. 5B; Fig. S5E**).

**Figure 5.**
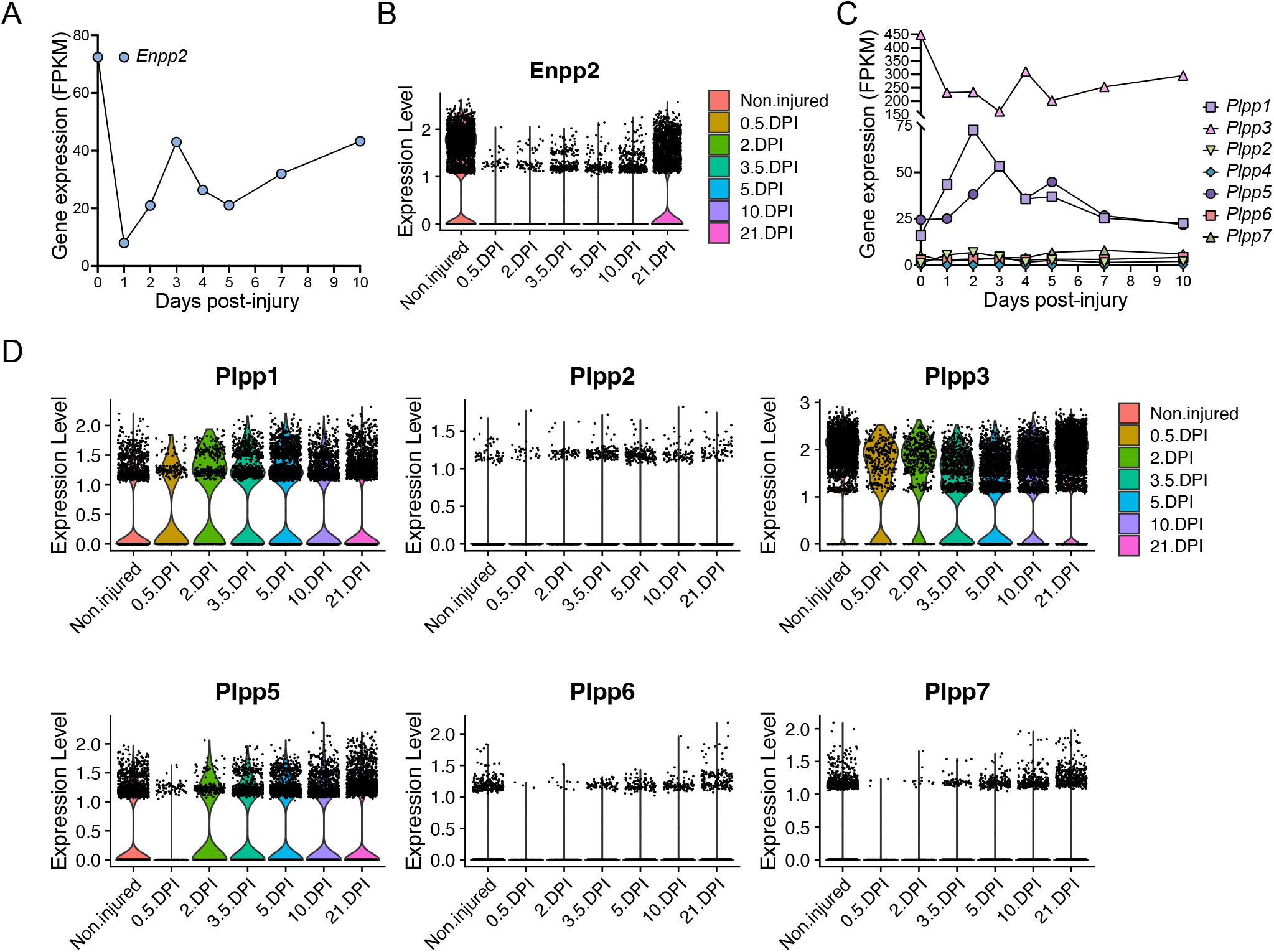
Quick and pronounced gene repression of LPA-producing and -catabolizing enzymes in fibro-adipogenic progenitors following muscle damage. (A) Quantification of Enpp2 transcript abundance (FPKM) in Hic1+ tdTomato expressing cells in response to injury. (B) Violin plots showing the gene expression level of *Enpp2* in FAPs in response to injury. DPI: Days post-injury. (C) Quantification of different Plpps transcript abundance (FPKM) in Hic1+ tdTomato expressing cells in response to injury. (D) Violin plots showing the expression level of the seven *Plpp* genes in FAPs in response to injury. DPI: Days post-injury.

*Hic1*-expressing cells repressed *Plpp3* expression immediately following injury, whereas *Plpp1* and *Plpp5* expression were transiently increased (**Fig. 5C**). These changes largely align with data derived from our single cell results of FAPs, however with some differences potentially accounted for by the pooling of FAP subsets. In pooled FAPs, all expressed *Plpp* genes were transiently downregulated early with expression recovering at later regenerative time points (**Fig. 5E**). Overall, the trend is towards an initial downregulation of transcripts for *Enpp2*, that produces extracellular LPA, and different *Plpp* genes, although *Plpp1* and *Plpp5* show different kinetics in *Hic1*-expressing cells (i.e. FAPs).

### Single cell cross-validation of the Enpp2-Lpar-Plpp axis in skeletal muscle cells

We next aimed to validate our previous single-cell transcriptomics findings using three public scRNAseq datasets (McKellar et al., 2021; Yang et al., 2022; Zhang et al., 2022). Using the dataset of Zhang and colleagues, we first observed that *Lpar1, Lpar4, Enpp2* and *Plpp3* were expressed by muscle FAPs, whereas tenocytes also expressed *Lpar1* and *Plpp3* (**Fig. S6A, B**). Again, FAPs did not express *Lpar2, Lpar3*, and *Lpar5* genes. *Lpar6, Plpp1*, and *Plpp3* were present in endothelial cells and pericytes (**Fig. S6A, B**). Some satellite cells express *Lpar1* and *Lpar4*, but not much of other LPA axis components (**Fig. S6A, B**). Neuron cells have a similar *Enpp2-Lpar-Plpp* expression profile to that of tendon cells (**Fig. S6A, B**). In addition, other two recently published skeletal muscle single-cell datasets further corroborated our previous findings (**Fig. S6C, D**) (McKellar et al., 2021; Yang et al., 2022). Of note, the dataset of Yang and colleagues used forelimb *triceps brachii* skeletal muscle, which supports our findings exploring the dataset of Oprescu et al. using hindlimb *tibialis anterior* muscle (Oprescu et al., 2020b; Yang et al., 2022). Furthermore, Yang and colleagues’ scRNAseq dataset from subcutaneous white adipose tissue also supports the notion of progenitor adipose stromal cells (ASCs), as cells highly expressing *Lpar1, Lpar4, Enpp2* and *Plpp3* (**Fig. S6E**), implying that the Enpp2-Lpar-Plpp axis is enriched in stromal cells from different tissue origins. These data, coupled with the observation that *Lpar1* and *Enpp2* are specific to skeletal muscle FAPs, supports a model where LPA could be involved in modulating the fate and behaviour of stromal cells, potentially through an autocrine signalling. Moreover, these gene expression profiles are consistent with our previous data analysis and conclusions, validating the dataset for further analysis.

### Extracellular LPA and LPAR-mediated downstream signalling are essential for fibro-adipogenic progenitor colony formation, growth, and proliferation

Given that the ENPP2-LPA-LPAR gene axis shows differential gene regulation in resting versus activated and proliferative subsets of FAPs, we hypothesized that the pathway could regulate fibro-adipogenic cell proliferation. Thus, we sought to evaluate FAP proliferation and cell cycle parameters in response to LPAR1 and LPAR3 subtype-selective antagonist Ki16425 (Ohta et al., 2003) and the potent ENPP2 inhibitor PF-8380 (Gierse et al., 2010) under conditions of colony formation and growth *in vitro*. FAPs have colony-forming units-fibroblast (CFU-Fs) properties, which reflects the presence of immature *in vivo* progenitors with proliferative, self-renewal and multi-lineage differentiation potential (Joe et al., 2010; Uezumi et al., 2010, 2014; Contreras et al., 2019b; Reggio et al., 2020; Farup et al., 2021). We evaluated the effects of Ki16425 and PF-8380 on SCA1^+^/PDGFRα^+^ FAP CFU-F formation and growth (**Fig. S7A**) and assessed colony numbers and self-renewal properties *in vitro* (**Fig. 6A**). Treatment of FAPs with Ki16425 significantly reduced FAP cell growth (**Fig. 6B**), suggesting that extracellular LPA, contained either in the bovine serum used for culture or endogenously produced by FAPs as they highly express *Enpp2*, has pro-proliferative effects. Consistently, PF-8380 treated cells formed only few colonies (**Fig. 6B**), suggesting that the LPA-producing activity of ENPP2 is essential for FAP proliferation and growth. We next utilized immunofluorescence and flow cytometric analyses to evaluate the percentage of DNA replicating cells, based on the incorporation of 5-ethynyl-2’-deoxyuridine (EdU) and its detection by click chemistry (Salic and Mitchison, 2008) (**Fig. 6C-F**). First, we evaluated the percentage of EdU^+^ FAPs at 24 h of inhibitor treatment in 10% FBS, after a short 2h pulse with EdU. Our data show that Ki16425 significantly reduced the proportion of cycling FAPs by half, as determined by the percentage of EdU^+^ cells (**Fig. 6D,E**). ENPP2 pharmacological inhibition with PF-8380 reduced the proportion of replicating FAP cells even more than Ki16425 inhibition of LPARs (**Fig. 6D,E**), corroborating our previous CFU-F findings. Quantitative detection of EdU^+^ FAPs using single-cell flow cytometry further corroborated our results (**Fig. 6F; Fig. S7B**). In addition, using Ki67 protein immune-labelling and flow cytometric detection, we show that Ki16425, and more profoundly PF-8380, significantly decreased the proportion of Ki67^+^ cycling-competent cells (**Fig. S7C**). LPA addition at 20μM did not rescue the reduction of cycling FAPs by Ki16425, and it only partially rescued the proliferation deficits induced by ENPP2 inhibitor (**Fig. 6F; Fig. S7C**), likely due to the presence of PLPPs. Further experiments showed that PF-8380 strongly blocks the progression of the G1-to-S phase transition of the FAP cell cycle (**Fig. 6F**), indicating that the ENPP2-LPA-LPAR axis regulates the cycling activity of FAPs.

**Figure 6.**
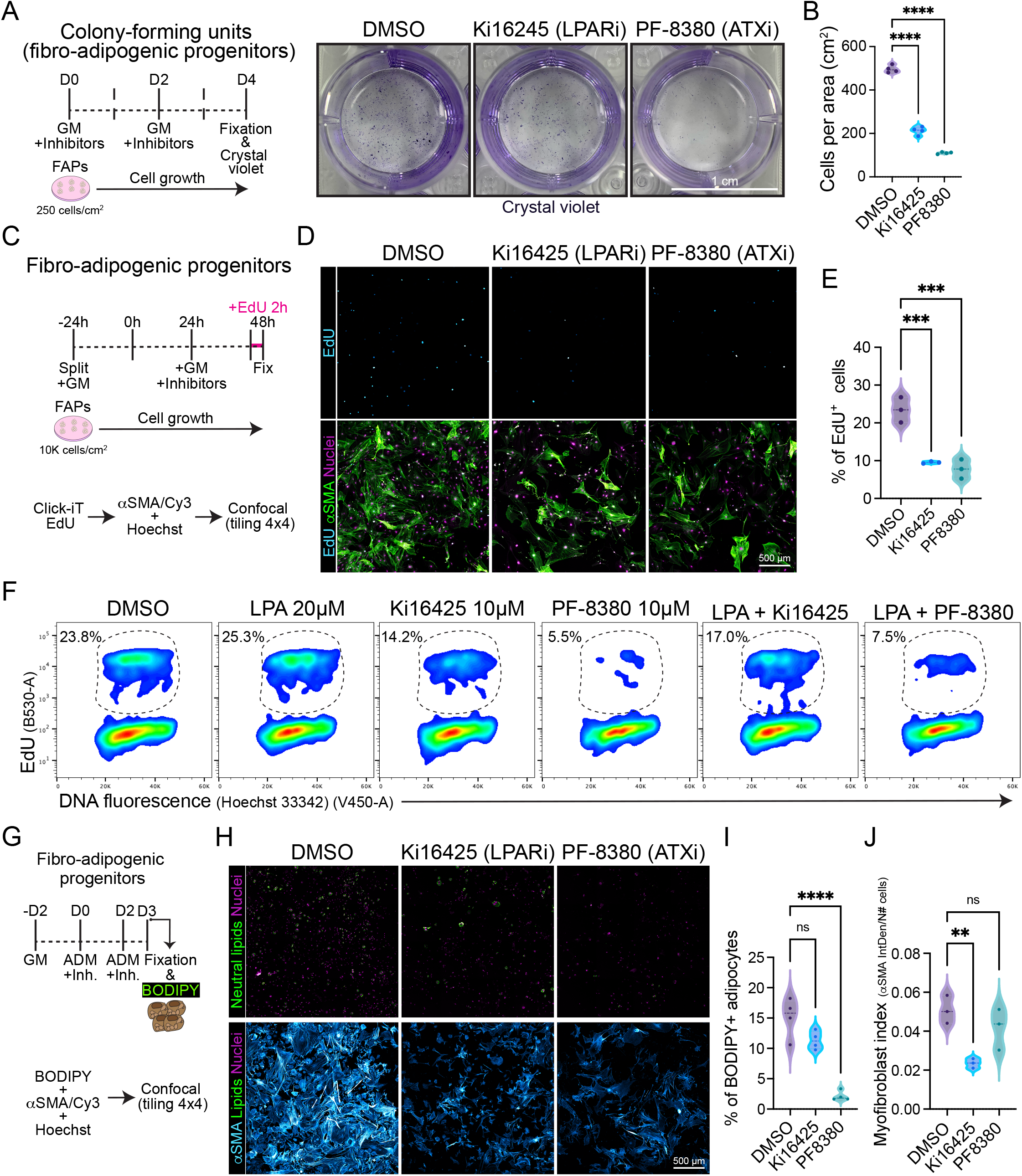
Pharmacological inhibiton of LPA receptors and Autotaxin reduces fibro-adipogenic progenitor cell growth and proliferation affecting their fibro-fatty fate. (A) Outline of colony-forming units assay using muscle FAPs. Representative images of FAPs control-treated (DMSO) or treated with Ki16245 (10μM, LPAR1/3 inhibitor) and PF-8380 (10μM, ATX inhibitor) as shown, and then stained with Crystal Violet. Scale bar: 1 cm. (B) Quantification of the number of cells per area as shown in (A) from four independent experiments. ****P < 0.0001 by one-way ANOVA with Tukey’s multiple comparison post-test; n = 4. (C) EdU assay outline. (D) Representative laser confocal images of FAPs after the different treatments (DMSO as control, Ki16245 (10μM) and PF-8380 (10μM)) at 24 h post treatment. EdU staining in shown in *cyan hot*, nuclear staining with Hoechst in *magenta*, and αSMA is shown in *light green*. Scale bar: 500 μm. (E) Quantification of the % of EdU labelled cells at 24 h post treatments. ***P < 0.001 by one-way ANOVA with Tukey’s post-test; n = 3. (F) Flow cytometry determination of EdU labelled cells in combination with DNA fluorescence at 24 h of treatments. (G) Outline of neutral lipid staining assay in muscle FAPs treated with adipogenic differentiation media (ADM). (H) Representative laser confocal images of FAPs after the different treatments (DMSO as control, Ki16245 (10μM) and PF-8380 (10μM)) at day 3 post treatment. BODIPY staining in shown in *light green*, nuclear staining in *magenta*, and αSMA in *cyan hot*,. Scale bar: 500 μm. (I) Quantification of the % of BODIPY labelled cells. ****P < 0.0001 by one-way ANOVA with Tukey’s multiple comparison post-test; n = 4. (J) Myofibroblast index of αSMA labelled cells. **P < 0.0021 by one-way ANOVA with Tukey’s multiple comparison post-test; n = 3.

Given that ENPP2 has been suggested as an adipose tissue-derived LPA generator, and we have shown that the *Enpp2-Lpar-Plpp* axis regulates skeletal muscle stromal FAP cell cycle and division, we next evaluated whether Ki16425 and PF-8380 could also impair cell growth and proliferation of visceral adipose stromal cells (ASCs) *ex vivo*. Our results show that both Ki16425 and PF-8380 inhibited the formation of ASC CFU-F (**Fig. S7D, E**). As found for FAP CFU-Fs, PF-8380-treated ASCs exhibited no cell growth (**Fig. S7D, E**). Both Ki16425 and PF-8380 treatments reduced the proportion of ASCs that can be detected in S-phase (i.e. EdU+), indicating defects on the G1-S phase transition (**Fig. S7F, G**). Flow cytometric analyses of DNA replicating cells indicated that LPA treatment (20 μM) without inhibitors induced a slight increase of EdU^+^ ASCs (**Fig. S7G**). LPA co-treatment with PF-8380 showed a small but significant rescue of the proportion of EdU^+^ ASCs compared to PF-8380 alone (**Fig. S7G**). Overall, these results support the notion that the ENPP2-LPA-LPAR axis regulates the proliferation and cell division of stromal cells from different tissue origins.

### Pharmacological inhibition of LPA1/3 receptors and ENPP2 impairs the differentiative fate of fibro-adipogenic progenitors

Finally, we evaluated the adipogenic differentiation of FAPs *in vitro*, scoring for FAP-derived adipocytes positive for neutral lipophilic molecule BODIPY staining at day 3 of induction using confocal tile image reconstruction (**Fig. 6G**). We observed that both LPAR and ENPP2 inhibition, from the beginning of differentiation, reduced the proportion of BODIPY^+^ adipocytes; however, only the more pronounced effect of ENPP2 inhibitor PF-8380 was statistically significant when normalised to total cell number (**Fig. 6H, I**). Both inhibitors also showed a strong anti-proliferative effect in adipogenic media, which reinforced our previous results (**Fig. 6H**). Since FAPs can also differentiate into activated fibroblasts and myofibroblasts, we evaluated alpha smooth muscle actin (αSMA)-positive stress fiber labelling as a proxy for myofibroblastic differentiation (**Fig. 6H**). Our results show that Ki16425 treatment leads to smaller sized FAPs and significantly reduced αSMA^+^ myofibroblast differentiation (**Fig. 6H, I**). However, whereas the ENPP2 inhibitor PF-8380 reduced myofibroblast differentiation of FAPs overall, this was not statistically significant when normalized to the total number of cells (**Fig. 6H, J**), highlighting the stronger anti-proliferative effect. Taken together, these results suggest that inhibition of LPA receptors and ENPP2 impairs the proliferative and fibro-fatty fate of fibro-adipogenic progenitors.

### Downregulation of *Lpar1* and *Lpar4* is associated with dividing and committed muscle stem cell state

Ray and colleagues recently reported that ENPP2-LPA-LPAR1 signalling is a crucial pro-regenerative axis in skeletal muscle (Ray et al., 2021). The authors also reported that *Lpar1* expression increased in myotubes compared to proliferative myoblasts, suggesting a pivotal role of LPA in modulating adult satellite cell differentiation.

To better understand the single-cell gene expression dynamics of the LPAR family in adult MuSCs, we again used scRNA-seq data and performed unsupervised sub-clustering on the MuSC metacluster, as previously shown (Oprescu et al., 2020; Contreras et al., 2021a). Six different subsets of MuSCs resulted from our analysis, consistent with previous findings (Oprescu et al., 2020; Contreras et al., 2021a) (**Fig. S8A**). Quiescent (QSC) adult MuSCs expressed *Lpar1* (about 50% of MuSCs) and *Lpar4* (20 % of MuSCs), but no other LPAR gene family members (**Fig. S8B**). *Lpar1* and *Lpar4* remained relatively stable in activated MuSCs (ASC), but decreased in dividing (DIV), committed (COM), immunomyoblasts (IMB) and differentiated (DIF) MuSCs (**Fig. S8B**), suggesting in fact that both LPA receptors are downregulated as muscle stem cells proliferate, commit, and differentiate to form mature myofibers.

Among phospholipid phosphatases expressed in quiescent MuSCs, *Plpp3* was the most highly expressed member of the family with expression progressively decreasing as these cells become activated, committed and differentiated (**Fig. S8B**). In contrast, *Plpp1* was significantly higher in activated, committed, and differentiating MuSCs, whereas *Plpp2* increased only in immunomyoblasts, and activated and dividing MuSCs (**Fig. S8B**). *Plpp5* and *Plpp6* were not detectably expressed in MuSCs (**Fig. S8B**). The non-enzymatic member, *Plpp7*, was absent in each of the six MuSCs subpopulations except for differentiated MuSCs (**Fig. S8B**). *Enpp2* expression was detected in ~20% percent of quiescent MuSC (**Fig. S8B**) and this further decreased in ASC, DIV, IMB, COM and DIF MuSCs subpopulations (**Fig. S8B**). Thus, our results suggest that *Enpp2* is expressed in at least some MuSCs, and functional data of Ray et al. suggest that this is sufficient to have biological relevance for muscle regeneration. These single cell transcriptomic analyses suggest that the expression of *Enpp2-Lpar-Plpp* axis genes is dynamic in muscle stem cells in homeostasis and injury. They illustrate also the potentially complex cell communication networks mediated by the bioactive phospholipid LPA in skeletal muscle and the MuSC niche.

### Pharmacological inhibition of ENPP2 inhibits satellite cell proliferation and myotube differentiation

Because our results show that *Lpar1* and *Lpar4* receptor genes are downregulated as muscle satellite cell proliferate, commit, and differentiate, we evaluated whether inhibiting LPAR1/3 receptors or ENPP2 would affect MuSC proliferation and differentiation. Our flow cytometric results showed that 10 μM of the ENPP2 inhibitor PF-8380 reduced by half the proportion of EdU^+^ satellite cells (**Fig. 7B, C; Fig. S9A**), as well as and the proportion of mitotic pH3^S10+^ cells (**Fig. 7D**). However, Ki16425 pharmacological inhibition of LPAR1/3 did not affect the proportion of EdU^+^ or pH3^S10+^ satellite cells (**Fig. 7C, D**), perhaps other receptors than LPAR1/3 may be involved. These data show that ENPP2 pharmacological inhibition impaired the number of replicating satellite cells. Next, we studied whether pharmacological inhibition of LPAR or ENPP2 would alter myotube differentiation of satellite cells. By evaluating myotube differentiation at day 3 (**Fig. 7E**), we observed that inhibition of LPAR with 10 μM Ki16425 did not affect myotube differentiation of satellite cells (**Fig. 7F**). By contrast, 10 μM of the ENPP2 inhibitor PF-8380 resulted in a significant reduction of sarcomeric α-ACTININ^+^ myotubes and MF20^+^ myotubes (**Fig. 7F; Fig. S9B**). Remarkably, we observed that 1 μM of PF-8380 also inhibited myotube differentiation and formation (**Fig. S9C**). Hence, ENPP2 catalytic activity is required for proper myotube differentiation and maturation, as previously suggested using a higher concentration of PF-8380 (25 μM) *ex vivo* (Ray et al., 2021). 20 μM LPA treatment alone significantly increased myotube (α-ACTININ^+^ and MF20^+^) number and thickness compared to untreated (control) cells (**Fig. 7F; Fig. S9B**). Overall, our data suggest that the LPA pathway is indispensable for myogenic differentiation of satellite cells.

**Figure 7.**
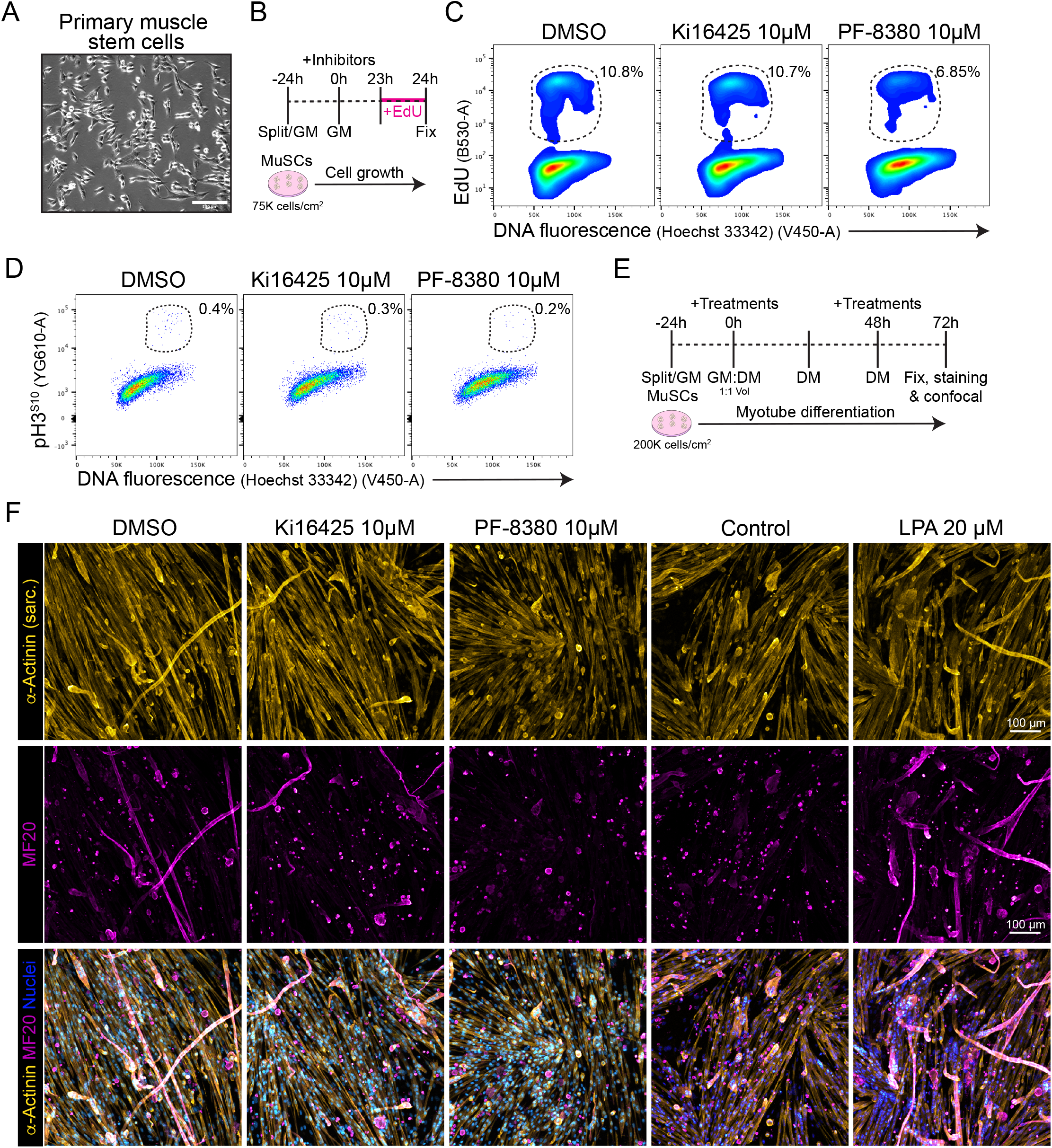
PF-8380 pharmacological inhibition of Autotaxin impairs satellite cell proliferation and myotube differentiation. (A) Brightfield images of cultured muscle stem cells (i.e. satellite cells). (B) Outline of EdU assay in muscle satellite cells. (C) Flow cytometry detection of EdU labelled cells in combination with DNA fluorescence at 24 h of treatments. (D) Flow cytometry detection of mitotic (phospho-H3^S10^) labelled cells in combination with DNA fluorescence at 24 h of treatments. (E) Outline of satellite cell differentiation protocol GM: growth media; DM: differentiation media. (F) Representative laser confocal images of day 3 myotubes after the different treatments. α-Actinin staining in shown in *hot yellow*, nuclear staining in *hot blue*, and MF20 in *magenta,*. Scale bar: 100 μm.

## Discussion

LPA is a signalling lipid with multiple biological functions and roles in health and disease. Collectively, our study provides insights into the presence and dynamic expression of the ENPP2-LPAR-PLPP gene axis in different muscle cells, cell lineages and states in homeostasis, injury and regeneration at single cell resolution. We first showed that *Lpar1* is the highest expressed member among other LPAR genes in *tibialis anterior* limb muscle, followed by *Lpar6* and *Lpar4*. We also found that *Lpar2, Lpar3*, and *Lpar5* were almost unexpressed. *Enpp2* is a relatively abundant gene in *tibialis anterior*, and several *Plpp* genes were also expressed, including *Plpp1, Plpp3*, and *Plpp7*. Thus, we report most ENPP2-LPAR-PLPP pathway gene components are present in healthy adult skeletal muscle in mice. FAPs highly express *Lpar1* and *Enpp2*, suggesting that stromal cells may be the primary source of extracellular LPA and LPA-mediated signalling and functions in the muscle niche. We additionally validated these findings utilising other scRNAseq datasets.

By sub-clustering stromal fibro-adipogenic progenitors (FAPs), we identified different subpopulations representing distinct cell states with robust LPAR and ENPP2 transcriptome signatures in homeostasis. *Lpar1* was expressed mainly by different subset of FAPs and tenocytes, whereas *Enpp2* was mostly expressed by resting FAPs. We also showed that tissue injury triggered transient repression of LPA receptors and *Enpp2*. Hence, uniquely activated FAP cell states are partly defined by a downregulation of *Lpar* and *Enpp2* gene expression. In addition, our *ex vivo* experiments indicate that the LPAR1/3 Ki16425 receptor antagonist and ENPP2 inhibitor PF-8380 impaired cell cycle progression and proliferation of muscle FAPs and visceral ASCs, although PF-8380 had a stronger effect. Since *Lpar3* is not expressed by resting or activated FAPs, we speculate that most Ki16425-driven effects are mediated by inhibition of LPAR1 in FAPs. Here, in PF-8380 treated cells, we also found decreased adipogenic differentiation of FAPs, in part due to the proliferative deficits induced by this potent ENPP2 inhibitor. On the contrary, although Ki16425 treatment did not significantly impair adipocyte differentiation of FAPs, it did reduce the proportion of αSMA+ myofibroblasts. Thus, pharmacological inhibition of LPAR1 and ENPP2 reduced the growth and proliferation of stromal cells, affecting their differentiation potential.

Finally, focusing on different MuSCs subtypes that emerge following acute damage we also observed differential ENPP2-LPAR-PLPP axis gene expression, although in general terms the axis was more lowly expressed compared to FAPs and tenocytes. Activated muscle stem cell states associate with *Lpar1* and *Lpar4* upregulation, suggesting that as satellite cells abandon their resting state, they increase the expression of LPARs, which in turn could bind niche-stored or damaged-released LPA, increasing the responsiveness of satellite cell lineage cells to local LPA. In this study, using pharmacological inhibition, we found that ENPP2 was essential for satellite cell proliferation and myotube differentiation. We also reported that exogenous LPA is sufficient to enhance the efficiency of satellite cell differentiation and myotube maturation, indicating that even low transcript abundance of LPA receptors in satellite cells is enough to elicit relatively strong cellular responses to extracellular LPA *ex vivo*. Related to this finding, Ray and colleagues recently showed that myogenic differentiation induces *Enpp2* expression, suggesting an increase in the extracellular abundance of ENPP2, and *Enpp2* knockdown reduced fusion and myotube differentiation (Ray et al., 2021). Remarkably, the cell-specific deletion of *Enpp2* in MuSCs impairs acute injury-induced muscle regeneration in mice, resulting in reduced muscle fiber caliber (Ray et al., 2021). These results were supported by utilizing the pharmacological ENPP2 inhibitor PF-8380, which also caused reduced muscle regeneration. Furthermore, *Enpp2* transgenic mice overexpressing circulating and extracellular ENPP2 levels, and expectedly increasing serum LPA, showed signs of muscle hypertrophy via ribosomal S5K signalling and accelerated recovery post-acute damage (i.e., faster muscle regeneration) (Ray et al., 2021). In support of this, intramuscular injections of both ENPP2 and LPA into healthy muscles resulted in muscle hypertrophy. These results provide significant data in favour of a pro-regenerative role of the ATX-LPA-LPAR axis in skeletal muscles. Additionally, inhibition of ATX using PF-8380 at 25 μM impaired satellite cell differentiation into myotubes but did not affect satellite cell proliferation using 5-bromo-2’-deoxyuridine (BrdU) uptake (Ray et al., 2021). In contrast, our results showed that 10 μM PF-8380 was sufficient to alter EdU uptake and the mitotic mark H3 phosphorylated in Serine10 in satellite cells *ex vivo*, suggesting that ENPP2 regulates MuSC proliferation. Lower concentrations of the ENPP2 inhibitor PF-8380 than that used by Ray et al., (2021) also impaired satellite cell myotube differentiation, supporting the notion that ENPP2 activity is key for proper skeletal myogenesis. Overall, future studies are needed to understand the role of LPA on skeletal myogenesis and muscle regeneration. However, due to the pleiotropic effects LPA might have on different cell types and cell states, addressing these questions on models of muscle damage remain challenging.

Kienesberger and colleagues showed that the ENPP2-LPA axis is involved in obesity-induced insulin resistance in muscles, affecting mitochondrial respiration in differentiated myotubes (D’Souza et al., 2018), validating the previously suggested key role of ENPP2-LPA axis in healthy and obese adipose tissue (Ferry et al., 2003; Boucher et al., 2005; Federico et al., 2012). The authors also showed that partial genetic reduction of ENPP2 levels ameliorated obesity and systemic insulin resistance in a high-fat diet mouse model (D’Souza et al., 2018). These results suggest that the ENPP2-LPA axis could contribute to the development of obesity-related disorders and tissue malfunction in metabolically altered states. Our analysis shows that adipose tissue stromal cells highly express Enpp2, and do respond to ENPP2 inhibitor PF-8380.

We and others have demonstrated that LPA induces the gene expression and protein levels of biologically active Connective Tissue Growth Factor (CTGF/CCN2) in C2C12 myoblasts (Vial et al., 2008; Riquelme-Guzmán et al., 2018). CTGF is a matricellular regulatory protein that modulates skeletal muscle repair, muscular dystrophy pathophysiology, and fibrosis (Morales et al., 2013; Petrosino et al., 2018, 2019; Rebolledo et al., 2019; Chen et al., 2020). LPA-mediated CTGF induction has been reported in different cell types, including embryonic and adult fibroblasts, mesothelial cells, in human and mouse models (Heusinger-Ribeiro et al., 2001; stortelers et al., 2008; Sakai et al., 2013). Remarkably, several ubiquitous signalling pathways mediate LPA-mediated CTGF induction in myogenic cells, including αvβ3 and αvβ5 Integrins, TGF-β receptor kinase activity, JNK, ECM components, and FAK (Cabello-Verrugio et al., 2011; Riquelme-Guzmán et al., 2018). These studies reveal an intricate network of signalling molecules that may tune LPA-driven responses in cells and tissues. Remarkably, Chen and colleagues showed that LPA, which has been previously identified to increase upon myocardial infarction (Chen et al., 2003), promotes proliferation and apoptosis of cardiac fibroblasts depending on its concentrations, suggesting that LPA has dual roles in fibroblasts (Chen et al., 2006). Our flow cytometry and imaging data, highlighting cell cycle and proliferation analyses, show that LPA does not cause fibro-adipogenic progenitor cell death but, on the contrary, supports a pro-proliferative role of the ENPP2/LPA/LPAR axis in muscle and adipose tissue-derived stromal cells.

Recently, Córdova-Casanova and colleagues showed that intramuscular injections of LPA induced CTGF protein levels and a few ECM proteins (Córdova-Casanova et al., 2022). Using LPAR inhibitor Ki16425 or LPAR1 knockout mice the authors observed an inhibition of these effects. They also showed increase muscle cellularity, i.e., total number of nuclei, and number of PDGFRα^+^ FAPs in response to LPA intramuscular injections. These results indicate LPA could have fibrotic-like properties *in vivo* in damaged muscles as previously suggested using cell culture (Vial et al., 2008; Riquelme-Guzmán et al., 2018) or *in vivo* models (Davies et al., 2017). Periodic intraperitoneal injections of LPA worsen the inflammatory milieu of rotator cuff tears (RCTs) in adult rats, increasing *Tnfa* and *Tgfb1* at six weeks post tear and the number of inflammatory cells within the affected muscles (Davies et al., 2017). Rotator cuff tears (RCTs) are a highly prevalent form of muscle trauma and tissue degeneration (Agha et al., 2021). Since severe intramuscular fibrosis and fatty infiltration are key morphological features of RCTs, significant research suggests FAPs as critical mediators of RCT onset, pathology, and progression (Theret et al., 2021). The authors also showed that enhanced systemic LPA worsens the fibrotic and adipogenic phenotype of RCTs (Davies et al., 2017). Thus, their study is the first of its kind to demonstrate a pro-fibrotic and pro-adipogenic role of systemic LPA in damaged muscles. Although forced intramuscular injections with LPA may not reflect proper physiological or pathophysiological conditions, the results of Córdova-Casanova et al. together with those of Davis et al., offer a new avenue to start exploring the relevance of LPA-mediated signalling pathways and their role in muscle disease and physio-pathophysiology.

Since FAPs are the main mediators of ectopic fibrous and fatty tissue, and because our results show resting FAPs highly express *Lpar1* and *Enpp2*, we speculate that LPA acts on these stromal cells early after muscle injury to promote cell proliferation and survival, therefore, resulting in a fibrotic and adipogenic phenotype in severely damaged muscles. Our functional analyses demonstrated that LPA regulates the proliferative status of FAPs and ASCs, impacting also the differentiative fate of these cells. Owing that FAPs highly express *Enpp2* and *Lpar1*, we propose that an autocrine LPA signalling regulates the activation and proliferation of FAPs. Due our analysis also showed that endothelial cells and several immune cell types, including monocytes, M2 Cxc3cr1^hi^ macrophages, and APCs express *Lpar6*, we could also consider that intramuscular or intraperitoneal injections of LPA target endothelial and immune cells. In this regard, LPA promotes the development of macrophages from monocytes through a mechanism that may involve PPARγ (Ray and Rai, 2017). Hence, our results suggest that injury-induced LPA could act on monocyte to promote their maturation and differentiation into macrophages via LPAR6.

The involvement of the ENPP2-LPA-LPAR axis in inflammation and fibrosis is not new and several studies have shown its crucial participation (Tager et al., 2008; Castelino et al., 2011; Gan et al., 2011; Sakai et al., 2013; Ohashi and Yamamoto, 2015; He et al., 2018; Ninou et al., 2018); however, the mechanisms and cellular targets have been underexplored. Several ongoing studies suggest the ENPP2-LPA-LPAR axis as a prognostic indicator of injury- or radiation-induced fibrosis (NCT05031065), with some studying the safety, tolerance, and effectiveness of orally available ENPP2 inhibitors (BBT-877: NCT03830125; GLPG1690 (ziritaxestat): NCT02738801 and NCT03798366) or LPAR antagonists (BMS-986020: NCT01766817 (Decato et al., 2022); BMS-986278: NCT04308681), as a means of reducing tissue fibrosis and improving organ function in different patients cohorts. Because stromal cells of mesenchymal origin, e.g., FAPs and ASCs, highly express *Lpar1, Enpp2*, and key *Plpp* members, upcoming research should focus on better understanding the role of LPA axis in muscle homeostasis, inflammation, fibrosis, repair, and regeneration. This understanding would potentially offer new druggable avenues for devastating muscle diseases like myopathies, severe muscle trauma, or neuromuscular disorders.

In summary, applying bulk and single cell transcriptomic data analyses we zoom in on skeletal muscle tissue at single cell resolution and provide for the first time a detailed view of the ENPP2-LPA-LPAR-PLPP axis for future insights in how to target LPA-driven signalling and functions. Furthermore, using *ex vivo* FAP and adipose stromal cell cultures and pharmacological inhibition of LPARs and ENPP2, we demonstrate, for the first time, that the ENPP2-LPA-LPAR axis regulates the cell cycle activity and proliferation of these cells. Hence, our data analysis highlights LPA signalling in different muscle cells and fibro-adipogenic progenitor lineages after muscle injury and provides an entry point for more profound research of the role of LPA signalling in homeostasis, inflammation, fibrosis, repair, and regeneration.

### Limitations of the study

This study has certain limitations. First, our transcriptomics analyses cannot address the protein levels of the ENPP2-LPA-LPAR-PLPP network, noting, however, there is currently a limitation of validated and working antibodies of axis components. Second, although we detected a downregulation of LPARs in FAPs in response to injury, we did not evaluate LPA receptor protein levels in FAPs upon injury-induced activation. The development of high-quality and validated LPAR antibodies should help answer these and related questions. Third, commonly used tissue disaggregation strategies, flow cytometry, and droplet-based scRNAseq does not efficiently capture certain cell types (e.g., adipocytes) because of their high propensity to rupture and buoyancy. Furthermore, large cells (e.g., myofibers, nerves, and adipocytes) do not effectively fit into a droplet and are often underrepresented in scRNAseq studies. Fourth, we have not evaluated or measured the effects of exogenous LPA or pharmacological inhibitors of LPARs or ENPP2 on modulating the fate of immune, tenocytes, or endothelial cells. Subsequent studies should also focus on understanding the influence and role of the ENPP2-LPA-LPAR-PLPP network and its effects on the fate of different muscle cells. Nevertheless, our study represents the first of its kind since in exploring the ENPP2-LPA-LPAR-PLPP network at single-cell resolution, and the proliferative and differentiated fate of fibro-adipogenic progenitors with altered LPA signalling.

## Materials and methods

### scRNA-seq data processing and analyses

We extracted the single-cell RNA sequencing data used in this paper from Gene Expression Omnibus (GEO; GSE138826) (https://www.ncbi.nlm.nih.gov/geo/query/acc.cgi?acc=GSE138826; GSE138826_expression_matrix.txt) (Oprescu et al., 2020). The preliminary analyses of processed scRNA-seq data were analysed using the Seurat suite (version 4.0.3) standard workflow in RStudio Version 1.2.5042 and R version 4.0.3. First, we applied initial quality control to Oprescu et al., 2020 dataset. We kept all the features (genes) expressed at least in 5 cells and cells with more than 200 genes detected. Otherwise, we filtered out the cells. Second, we verified nUMIs_RNA (> 200 & < 4000) and percent.mt. (less than 5%) Third, UMIs were normalized to counts-per-ten-thousand log-transformed (normalization.method = LogNormalize). The log-normalized data were then used to find variable genes (*FindVariableFeatures*) and scaled (*ScaleData*). Finally, *Principal Component Analysis* (PCA) was run (*RunPCA*) on the scaled data using highly variable features or genes. *Elbowplot* were used to decide the number of principal components (PCs) to use for unsupervised graph-based clustering and dimensional reduction plot (UMAP) embedding of all cells or further subclustering analyses (i.e., FAPs) using the standard *FindNeighbors, FindClusters*, and *RunUMAP* workflow. We used 30 PCs and a resolution of 0.6 to visualize a Uniform manifold approximation and projection (UMAP) dimensionality reduction plot generated on the same set of PCs used for clustering. We decided the resolution value for FindClusters on a supervised basis after considering clustering output from a range of resolutions (0.4, 0.6, 0.8, and 1.2). We used a resolution of 0.6. Our initial clustering analysis returned 29 clusters (clusters 0-28). We identified cell populations and lineage-specific marker genes for the analysed dataset using the *FindAllMarkers* function with logfc.threshold = 0.25, test.use = “wilcox”, and max.cells.per.ident = 1000. We then plotted the top 10 expressed genes, grouped by orig.ident and seurat_clusters using the *DoHeatmap* function. We determine cell lineages and cell types based on the expression of canonical genes. We also inspected the clusters (in Figure 2 and Figure 3) for hybrid or not well-defined gene expression signatures. Clusters that had similar canonical marker gene expression patterns were merged.

For Mesenchymal Clusters (group of FAPs + DiffFibroblasts + Tenocytes obtained in **Figure 2**) we used PCs 20 and a resolution of 20 to visualize on the UMAP plot. Our mesenchymal subclustering analysis returned 10 clusters (clusters 0-9). Cell populations and lineage-specific marker genes were identified for the analysed dataset using the *FindAllMarkers* function with logfc.threshold = 0.25 and and max.cells.per.ident = 1000. We then plotted the top 8 expressed genes, grouped by orig.ident and seurat_clusters using the *DoHeatmap* function. The identity of the returned cell clusters was then annotated based on known marker genes (see details about cell type and cell lineage definitions in the main text, results section). Individual cell clusters were grouped to represent cell lineages and types better. Finally, figures were generated using *Seurat* and *ggplot2* R packages. We also used dot plots because they reveal gross differences in expression patterns across different cell types and highlight moderately or highly expressed genes.

To validate our initial skeletal muscle single-cell analysis, we explored three publicly available scRNAseq datasets (McKellar et al., 2021; Yang et al., 2022; Zhang et al., 2022). Zhang et al. dataset was explored using R/ShinyApp (https://mayoxz.shinyapps.io/Muscle), McKellar et al., 2021 using their web tool developed http://scmuscle.bme.cornell.edu/, and Yang et al. using their Single Cell Metab Browser http://scmetab.mit.edu/. All the figures used were downloaded from the websites (**Fig. S6**).

The scRNAseq pipeline used for MuSC subclustering was developed following previous studies (Oprescu et al., 2020; Contreras et al., 2021a). To perform unsupervised MuSC subclustering, we used Seurat’s subset function *FindClusters*, followed by dimensionality reduction and UMAP visualization (*DimPlot*) in Seurat.

### Bulk RNA-seq data processing and analyses

Bulk RNA-seq data was extracted as FPKM values from a previously processed dataset extracted from GEO (GSE110038) (Scott et al., 2019). No further RNA-seq processing was performed to that of Scott and colleagues. We generated the heat maps shown in **Fig. 1** and **Fig. 2** with Morpheus (https://software.broadinstitute.org/morpheus/) using previous transcriptomic available RNA-seq data (Scott et al., 2019).

### Reagents

We used oleoyl-L-α-lysophosphatidic acid sodium salt, LPA (L7260-1MG, Sigma-Aldrich), Ki16425 (potent antagonist of the lysophosphatidic acid receptors LPA1 and LPA3, SML0971-5MG, Sigma-Aldrich), PF-8380 (Autotaxin inhibitor, Cat. No. HY-13344, MedChemExpress). LPA was reconstituted according to the supplier’s instructions. Ki16425 and PF-8380 were reconstituted in cell culture grade Dimethyl sulfoxide (Hybri-Max DMSO, D2650, Sigma-Aldrich) at 10 mM stock according to the supplier’s instructions and used as indicated in the corresponding figures. DMSO was used as a control when these inhibitors were added. Ki16425 and PF-8380 were added at 15 min prior being co-incubated with LPA, when indicated. Other reagents, unless otherwise is indicated, were purchased from Sigma-Aldrich.

### Ethics Statement

All experimental procedures were approved by the Garvan Institute/St. Vincent’s Hospital Animal Experimentation Ethics Committee (No. 13/02, 16/10, 19/14) and performed in strict accordance with the National Health and Medical Research Council (NHMRC) of Australia Guidelines on Animal Experimentation. All efforts were made to minimize suffering.

### Mice

Wild type mice (Inbred C57BL/6J, Stock No: 000664, Jackson Laboratory) were bred and housed in the BioCORE facility of the Victor Chang Cardiac Institute. Rooms were temperature and light/dark cycle controlled, and standard food was provided ad libitum. Two-to four-month-old female mice were used in experiments regarding ex vivo culture of fibro-adipogenic progenitors and satellite cells.

### Skeletal muscle fibro-adipogenic progenitors and muscle stem cell isolation, *ex vivo* culture, and FAP CFU-F

One-step digestion of skeletal muscle tissue for fibro-adipogenic progenitor isolation was performed as described before with few modifications (Contreras et al., 2020). Briefly, skeletal muscles from both hindlimbs of female wild type mice were carefully dissected, washed with ice-cold DMEM, and cut into small pieces with blades until a homogeneous, paste-like slurry was formed. Seven ml of digestion solution containing collagenase type II (265 Unit/ml, Worthington, US), 0.5 U of Dispase (Cat. No. 07913, STEMCELL™ Technologies, Canada), 0.05 mg/mL of DNaseI (Cat. No. 10104159001, Roche/Sigma-Aldrich, 100mg from bovine), and 1% BSA (Sigma-Aldrich Pty Ltd, A3311-50G) dissolved in DMEM (Cat. No. 10566016) was added to two hindlimbs and the preparation was placed on a water bath with constant rotation at 37°C for 45 min and intermittent vortexing every 15 min. Tissue preparations were gently pipetted up and down 5-10 times to enhance muscle dissociation with a 10 mL Stripette^®^ serological pipette on ice. Ice-cold FACS buffer was added to make the final volume up to 30 mL volume and samples were then passed through 100 μm cell strainer sequentially after gentle mixing. Following centrifugation at 600 g for 6 min at 4C, the pellet was resuspended in 10 mL of growth media (20 ng/mL of basic Fibroblast Growth Factor (Milteny Biotec, Cat. No. 130-093-843) and 10% heat-inactivated fetal bovine serum (v/v) (FBS; Hyclone, UT, USA) in DMEM (Cat. No. 10566016) and supplemented with antibiotics (Penicillin-Streptomycin Cat. no. 15140122, Gibco by Life Technologies)) and cells were pre-plated onto 100 mm plastic tissue culture dish for 2 h and grown at 37°C in 5% CO_2_ as previously described (Contreras et al., 2019c). After 2h of FAP pre-plating the supernatant media was removed to culture muscle stem cells (see *Muscle stem cell enrichment and myotube differentiation* protocol below) and replaced with fresh growth media. FAP CFU-F assay was performed with cells seeded at a density of 250/cm^2^ in growth media in a 12-well plate coated with Corning Matrigel Matrix hESC qualified (Cat. No. 354277) prepared in cold DMEM/F-12 as per the provider’s instructions. Cultured FAPs were allowed to grow for about 7 days before splitting them. CFU-F experimental outline is shown in the figure. FAPs were used not further than passage 1. CFU-F averages were obtained from 3 technical replicates/samples using three female mice. CFU-F photos were taken using an iPhone XR 12MP Wide camera.

### Muscle stem cell enrichment and myotube differentiation

After 2 h of fibro-adipogenic progenitors pre-plating (as described above), muscle stem cells were enriched by transferring the muscle preparation supernatant into a new 100 mm plastic tissue culture dish coated with Corning Matrigel Matrix (as described above) and further cultured for 2 h. Then, the supernatant was carefully replaced with 10 mL of MuSC growth media (20 ng/mL of basic Fibroblast Growth Factor (Milteny Biotec, Cat. No. 130-093-843) and 10% heat-inactivated fetal bovine serum (v/v) (FBS; Hyclone, UT, USA) in DMEM (Cat. No. 10566016)). The MuSC growth media was replaced every second day and the cells were allowed to growth for 4-5 days before splitting them. Muscle stem cells were used not further than passage 1. MuSCs were platted at 75,000 cells per cm^2^ when EdU (at 10 μM final concentration) or pH3^S10^ labelling (Alexa Fluor^®^ 594 anti-Histone H3 Phospho (Ser10) Antibody, 1:250 dilution, clone 11D8, Cat. No. 650810, Biolegend) was performed as indicated in **Fig. 7B-D**. Hoechst 33342 was used at 10 μg/ml final concentration. For MuSC-into-myotube differentiation, MuSCs at passage 0 were split using MuSC growth media at 200,000 cells per cm^2^ and cultured into 48-well plates coated with hESC-qualified Corning Matrigel Matrix for 24 h. Then, 500 μL of myotube differentiation media (5% of Horse serum (H1270-100ML, Sigma-Aldrich) in DMEM (Cat. No. 10566016)) were added to 500 μl of MuSC growth media. The mixed media was then changed every day using myotube differentiation media. Cells were fixed in 4% PFA for 15 min and kept in PBS1x at 4C until myotube staining was performed. Myotubes were permeabilized in 1× saponin-based permeabilization and wash buffer (0.2% (w/v) saponin containing 4% (v/v) FBS (v/v), 1% (w/v) BSA and 0.02% (v/v) Sodium Azide in PBS) for 10 min. Cells were then stained for 2 h using sarcomeric α-Actinin antibody (α-Actinin (Sarcomeric) Antibody, anti-human/mouse/rat, Vio^®^ R667, REAfinity™, clone REA402, 130-128-698, 1:100 dilution) and MF20 (MF 20 was deposited to the DSHB by Fischman, D.A. (DSHB Hybridoma Product MF 20), 1:20 dilution, Uniprot ID: P13538 (Myosin heavy chain, sarcomere (MHC)) and Hoechst 33342 at 10 μg/ml. Confocal laser scanning microscopy of stained myotubes was performed using a LSM900 Inverted confocal laser scanning microscope that comprises an upright Zeiss Axio Observer 7, four laser lines, two Gallium Arsenide Phosphide photomultiplier tubes (GaAsP-PMT), and a motorised stage. 4×4 tile images were acquired on a Zeiss Axio Observer 7 fitted with an LSM 900 confocal scan head, using a 10X objective, 0.45 numerical aperture with a z-step size of 4 μm, 1024 x 1024 μm, WD 2.0, Plan-APO UV-VIS–NIR, and PBS immersion.

### Flow Cytometry of fibro-adipogenic progenitors using stromal markers

Flow cytometry analyses of FAP markers were performed in day 6-7 growing FAPs at passage 0 at 70–80% confluence using a BD LSRFortessa™ X-20 Cell Analyzer. We used freshly TrypLE-dissociated FAPs. Briefly, FAPs were dissociated in 1 ml (6-well plate) of TrypLE as described before. After TrypLE incubation, 0.5 ml of cold FACS Buffer was added, cells were fully dissociated, samples collected in 2 ml tubes and centrifuged at 500×g for 5 min. The supernatant was carefully discarded, and the pellet of cells resuspended thoroughly with 1 ml ul of cold FACS Buffer. Total protein labelling (**Fig. S7A**) was determined by flow cytometry through fixing the dissociated cells in 2% PFA for 10 min at 4°C. After fixation, cells were washed three times, with 3 ml of PBS 1×. Cells were stained with primary antibodies for 30 min at RT in in BD perm/wash buffer at ~2.5 × 10^5^ cells per 100 μL of cell suspension. The following antibodies were used: FITC Rat Anti-Mouse Ly-6A/E (SCA-1) (Clone E13-161.7), BD Bioscience (1:200 dilution, Cat. no. 561077), PE anti-mouse CD140a (PDGFRA)

Monoclonal Antibody (APA5), BioLegend (1:200 dilution, Cat. no. 135905, Lot: B244566), APC anti-mouse CD140b Rat IgG2a, κ Antibody (APB5), BioLegend (1:200 dilution, Cat. no. 136007, Lot: B306888) and APC/Cyanine7 anti-mouse CD90.2 Rat IgG2b, κ Antibody, BioLegend (Clone 30-H12) (1:200 dilution, Cat. No. 105327, Lot: B353527). The isotypes control antibodies used were as follows: FITC Rat IgG2a, κ Isotype Control (Clone R35-95, BD Bioscience, Cat. no. 553929), PE Rat IgG2a, κ Isotype Ctrl Antibody (BioLegend, Clone RTK2758, Cat. no. 400507), APC Rat IgG2a, κ Isotype Ctrl Antibody (BioLegend, Clone RTK2758, Cat. no. 400511), and APC/Cyanine7 Rat IgG2b, κ Isotype Ctrl Antibody (BioLegend, Clone RTK4530, Cat. no. 400623). After staining, cells were washed three times after staining using BD perm/wash buffer, and analyzed by flow cytometry. All flow cytometry data were analyzed using FlowJo software (version 10.8.1, BD).

### Adipogenic differentiation of fibro-adipogenic progenitors and adipocyte assessment

After 6-8 days of cell growth, passage 0 FAPs were dissociated in 2 ml (100 mm culture dish) of pre-warmed TrypLE for 10 minutes. 10,000 FAPs per cm^2^ were added into each well using a 48-well plate and cells were allowed to grow for an additional of 1 up to 2 days using FAP growth media until the cells reached 95-100% confluence. Adipogenic differentiation was induced for 3 days using an in-house adipogenic induction media (ADM; 5% FBS, 1XPenStrep, 1 μM Dexamethasone, 0.5 mM IBMX, 1 μg/mL Human Insulin and 1 μM Rosiglitazone in high-glucose DMEM + GlutaMax). Then, cells were fixed in 4% PFA for 10 min at room temperature, washed with PBS and permeabilized in 1× saponin-based permeabilization and wash buffer (0.2% (w/v) saponin containing 4% (v/v) FBS (v/v), 1% (w/v) BSA and 0.02% (v/v) Sodium Azide in PBS) for 10 min. Cells were incubated for 1 h with 200 nM of BODIPY 493/503 (Cat.No. 25892, Cayman Chemical), αSMA-Cy3™ (1:200 dilution, clone 1A4, Cat. No. C6198, Sigma-Aldrich) and Hoechst 33342 (10 μg/ml in PBS, B2261-25MG, Sigma-Aldrich) in permeabilization and wash buffer at room temperature. Images were acquired on a LSM900 confocal laser microscope as detailed before. In brief, 4×4 tiled images were acquired (2.65 mm^2^ area at the center point of the well) and the total cell number and the percentage of BODIPY+ cells were quantified using Fiji software. Total cell number was determined using StarDist 2D plugin using the nuclei Hoechst staining layer. BODIPY+ adipocytes were counted manually using the Cell Counter plugin, and the values expressed as the % of BODIPY+ cells. Myofibroblast index was calculated quantifying the fluorescence intensity of αSMA-Cy3, normalized by the total number of cells per area.

### Cell cycle S-phase analysis of fibro-adipogenic progenitors and muscle stem cells using Click-iT EdU flow cytometry assay

5-ethynyl-2 deoxyuridine (EdU) flow cytometry analysis was determined in fibro-adipogenic progenitors and MuSCs as previously described (Contreras et al., 2021a). Briefly, 22 h after DMSO, LPA 20 μM or LPAR1/3 (Ki16425) or ATX (PF-8380) inhibitors treatments, EdU (10 μM final concentration) was added to the culture medium and incubated for 2 h. For negative staining controls, we included DMSO-treated cells that have not been exposed to EdU. Once each experimental condition and treatment was finished, cells were washed with PBS and dissociated in 1 ml (6-well plate) of pre-warmed TrypLE (TrypLE™ Express Enzyme (1×), no phenol red, Cat. no. 12604013, ThermoFisher Scientific). TrypLE was incubated for 7 up to 10 min at 37°C. After TrypLE incubation, 0.5 ml of cold FACS Buffer (PBS 1×, 2% FBS v/v, 2 mM EDTA pH 7.9) was added, cells were fully dissociated by pipetting, and samples collected in 2 ml tubes. Samples were centrifuged at 500 ×g for 5 min. Then, the supernatant was discarded, and the pellet of cells resuspended with 0.4 ml of cold PBS 1×. Cells were fixed by adding 0.5 ml of 4% PFA into ~0.5 ml of cell suspension. Cells were incubated for 10 min at room temperature, and then 2% PFA was washed three times with abundant PBS. When cells were ready to work with, they were distributed into 1.5 ml tubes, 500 μL of 0.1% BSA in PBS added, and pelleted. Pelleted cells were flicked and 400 μl of a 1× saponin-based permeabilization and wash reagent (0.2% (w/v) saponin containing 4% (v/v) FBS (v/v), 1% (w/v) BSA and 0.02% (v/v) Sodium Azide in PBS) was added and incubated for 15 min. Then the cells were centrifuged (500 ×g for 5 min). After the last centrifuge, EdU detection was performed using an in-house developed Click-iT EdU reaction cocktail made of 200 nM AZDye™ 488 Azide (Cat. No. 1275, Click Chemistry Tools, Scottsdale, AZ), 800 μM Copper (II) sulfate, and 5 mM Ascorbic acid in PBS1x. In brief, 400 μl per sample of the Click-iT reaction cocktail was added to the pellet, and the cells resuspended and incubated for 45 min at room temperature, protected from light. After Click-iT EdU reaction cocktail incubation, the cells were washed twice with 0.5 ml of 1× saponin-based permeabilization and wash reagent and pelleted at 500×g for 5 min, leaving 50 μl of pellet per tube which was resuspended by flicking. Then, 50 μl of conjugated antibodies prepared in perm/wash buffer (Ki-67 Antibody, anti-human/mouse, Vio^®^ R667, REAfinity™, order no. 130-120-562, clone REA183, 1:100 dilution, Miltenyi Biotech) and/or (Alexa Fluor^®^594 anti-Histone H3 Phospho (Ser10), clone 11D8, Mouse IgG2b, κ, 1:250 dilution, BioLegend) were added and incubated at RT for 1h. After antibody incubation, 800 μl of 0.1% BSA in PBS was added to each tube, and samples were spun down at 500g for 6 mins. The supernatants were removed and 50 μl pellets resuspended by flicking and incubated with 300 μl of Hoechst 33342 (10 μg/ml final concentration, B2261-25MG, Sigma-Aldrich) for 10 min at room temperature in 1× saponin-based permeabilization and wash reagent. Samples were analyzed by flow cytometry for DNA content and EdU labelled cells using a BD LSR Fortessa Laser Cell Analyser (BD Biosciences, Erembodegem, Belgium) equipped with 5 excitation lasers (UV 355nm, Violet 405 nm, Blue 488 nm, Yellow/Green 561 nm, and Red 633 nm). EdU-AZDye™ 488 Azide, Ki67-Vio^®^ R667, and pH3^S10^-Alexa Fluor^®^594 fluorescence were detected with logarithmic amplification using the B530 (530/30), R670 (670/14), and YF610 (610/20), detectors, respectively, whereas Hoechst fluorescence was detected with linear amplification using the V450 (V450/50) detector. Flow cytometry measurements were run at a mid-flow rate, and the core stream allowed to stabilize for 5 s prior acquisition. Data were collected using FacsDIVA 8 software. For optimal Hoechst signal detection and cell cycle progression analyses, an event concentration of <800 events/s was used, and 20,000 events were captured. All flow cytometry data were analyzed using FlowJo Portal (version 10.8.1, Becton Dickinson & Company (BD)) using Mac OS X operating system.

### Statistical analysis

Mean±s.e.m. values, as well as the number of experiments performed, are indicated in each figure. Bulk RNAseq data were collected in Microsoft Excel, and statistical analyses were performed using GraphPad Prism 9.4.0 software for macOS Monterey. All bulk RNAseq datasets used to determine gene expression were analyzed for normal distribution using the Shapiro-Wilk test with a significant level set at alpha = 0.05. Statistical significance of the differences between the means was evaluated using the One-Way ANOVA test followed by post-hoc Dunnett or Tukey’s multiple comparisons tests, and the significance level was set at *P* < 0.05 (95% confidence interval). *P*-value style: GP: 0.0332 (*), 0.0021 (**), 0.0002 (***), < 0.0001 (****).

## Abbreviations

ASCs: Adipose stromal cells
ATX: Autotaxin
CTGF/CCN2: Connective tissue growth factor
ENPP2: Ectonucleotide pyrophosphatase/phosphodiesterase 2
FAPs: Fibro-adipogenic progenitors
GPCRs: G protein-coupled receptors
LPARs: LPA receptors
LPA: Lysophosphatidic acid
MSCs: Mesenchymal stromal cells
MuSCs: Muscle stem cells
PA: Phosphatidic acid
PLPPs: Phospholipid phosphatases
scRNAseq: Single-cell RNA sequencing

## Data accessibility

This article has no additional data. Software used to analyze the data freely or commercially available.

## CRediT Author Contribution

**Osvaldo Contreras**: Conceptualization, Data curation, Formal Analysis, Investigation, Methodology, Project administration, Resources, Software, Supervision, Validation, Visualization, Writing – original draft, Writing – review & editing. **Richard P Harvey**: Resources, Supervision, Visualization, Review & Editing. The authors gave final approval for publication and agreed to be held accountable for the work performed therein.

## Acknowledgements

The authors would like to thank Drs Yen Tran, Qing Wang, and Camilo Riquelme-Guzmán for their help proofreading this manuscript, and Chris Thekkedam for his technical assistance. The authors also thank the Victor Chang Cardiac Research Institute Innovation Centre (Micro Imaging Facility) for technical support in image acquisition.

