## Supplemental Figure 1 for "Single-Cell Transcriptome Dynamics of the Autotaxin-Lysophosphatidic Acid Axis During Muscle Regeneration Reveal Proliferative Effects in Mesenchymal Fibro-Adipogenic Progenitors"

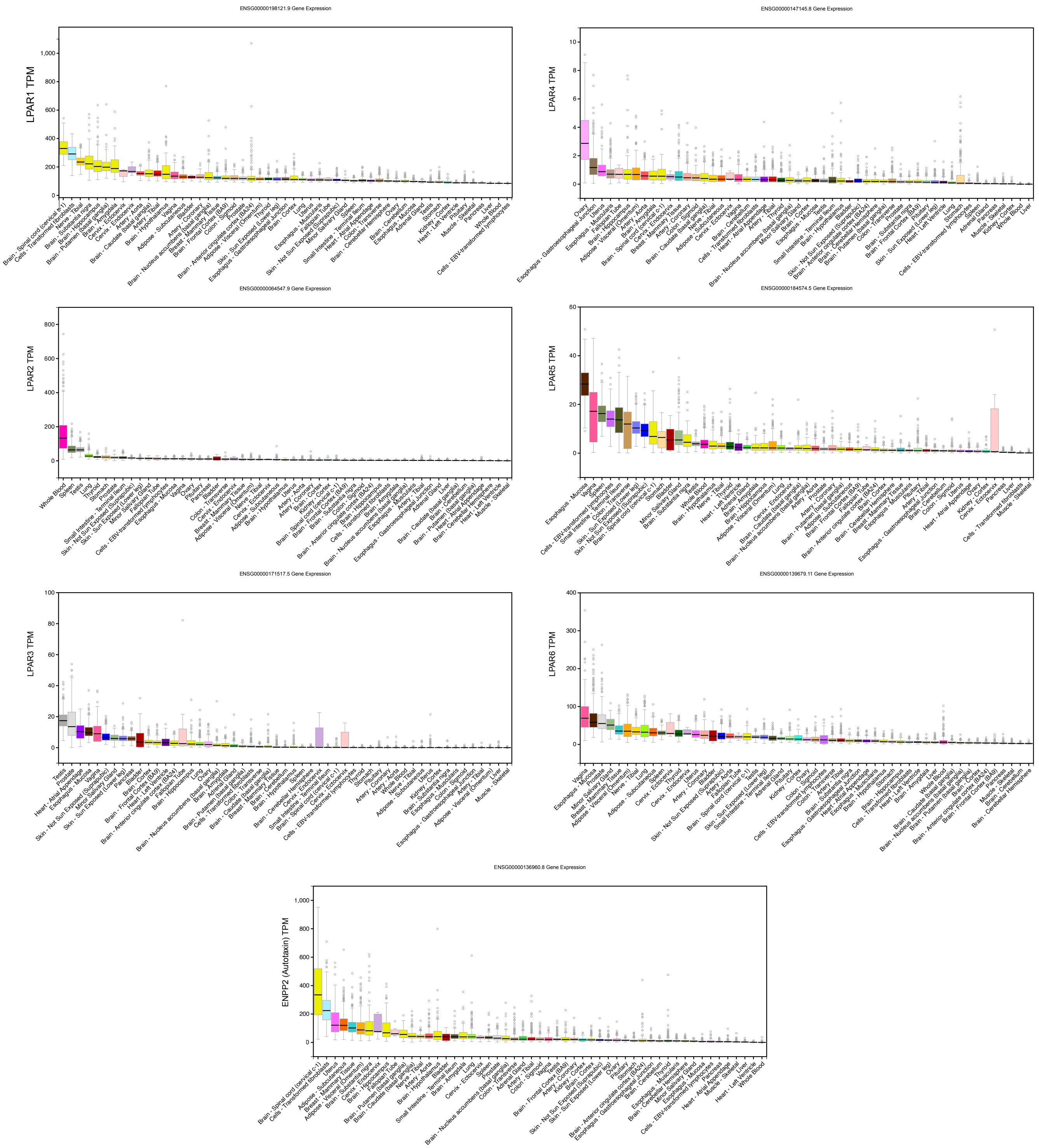


**Supplementary figure 1. Bulk tissue gene expression dynamics of the LPAR-Enpp2/Atx axis using GTEx portal.** Plots showing gene expression levels of LPAR and Enpp2 genes in different tissues and cells. Gene expression shown in transcript per million (TPM) linear scale. Bulk gene expression was sorted from high to low median expression. Data was downloaded from https://www.gtexportal.org/.
