## Supplemental Figure 2 for "Single-Cell Transcriptome Dynamics of the Autotaxin-Lysophosphatidic Acid Axis During Muscle Regeneration Reveal Proliferative Effects in Mesenchymal Fibro-Adipogenic Progenitors"

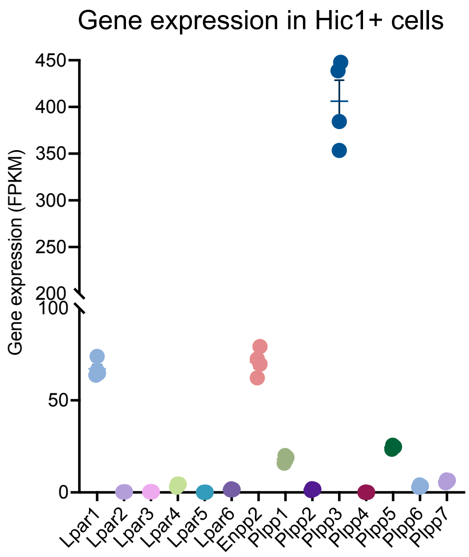


**Supplementary figure 2. Differential gene expression of the LPAR-Autotaxin-Plpp network in mesenchymal stromal cells.** (A) Quantification of Lpar, Enpp2, and Plpp transcript abundance (FPKM) in Hic1+ tdTomato expressing cells [Scott et al., 2019].
