## Supplemental Figure 3 for "Single-Cell Transcriptome Dynamics of the Autotaxin-Lysophosphatidic Acid Axis During Muscle Regeneration Reveal Proliferative Effects in Mesenchymal Fibro-Adipogenic Progenitors"

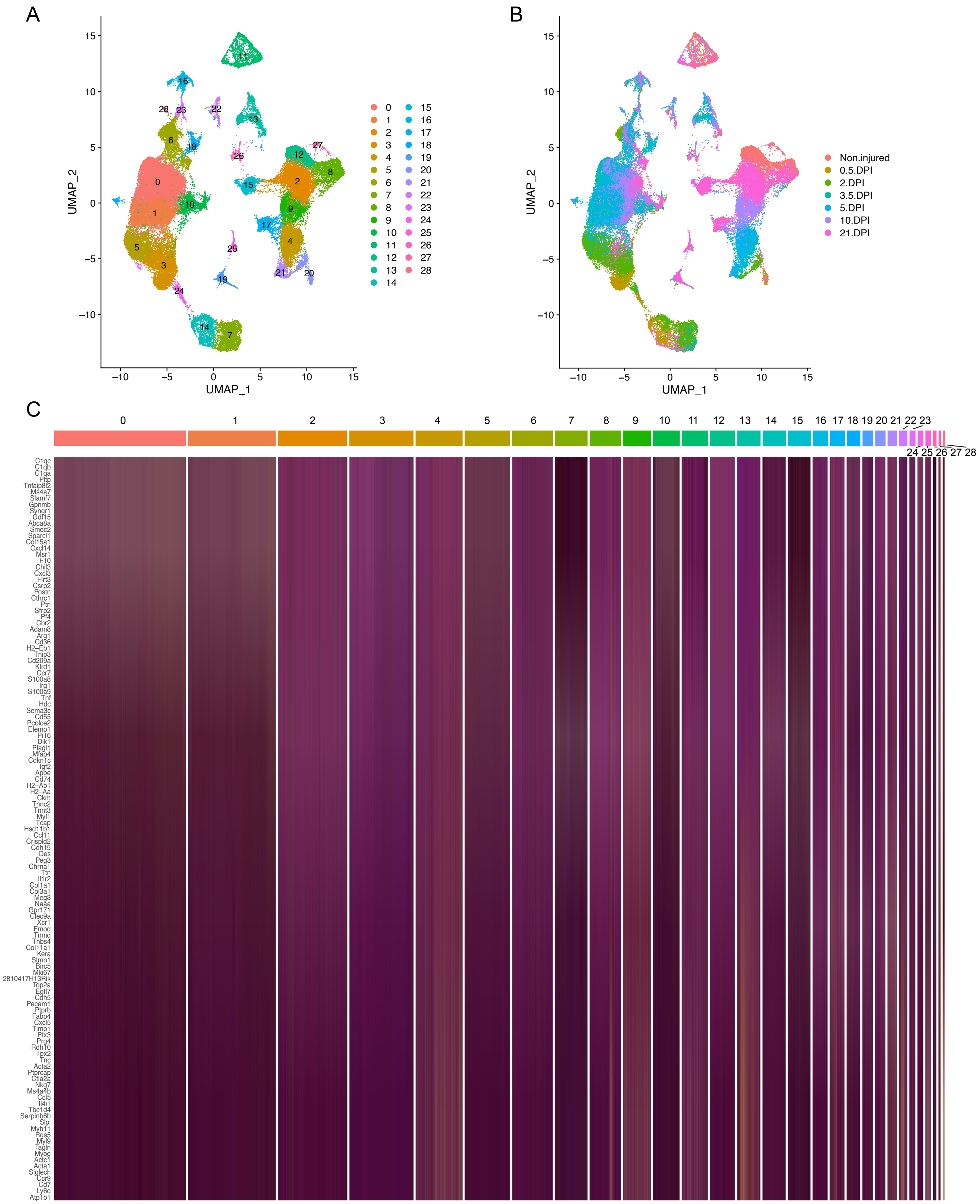


**Supplementary figure 3. Initial scRNAseq seurat-based clustering analysis and conditions.** (A) UMAP plot showing 29 distinct clusters across skeletal muscle homeostasis and regeneration. (B) UMAP plot showing distinct clusters based in non-injured or injured conditions. (C) Heat map plot of the 29 clusters shown in (A) and ordered based on the number of cells and the *top 5* expressed genes in each cluster subset.
