## Supplemental Figure 4 for "Single-Cell Transcriptome Dynamics of the Autotaxin-Lysophosphatidic Acid Axis During Muscle Regeneration Reveal Proliferative Effects in Mesenchymal Fibro-Adipogenic Progenitors"

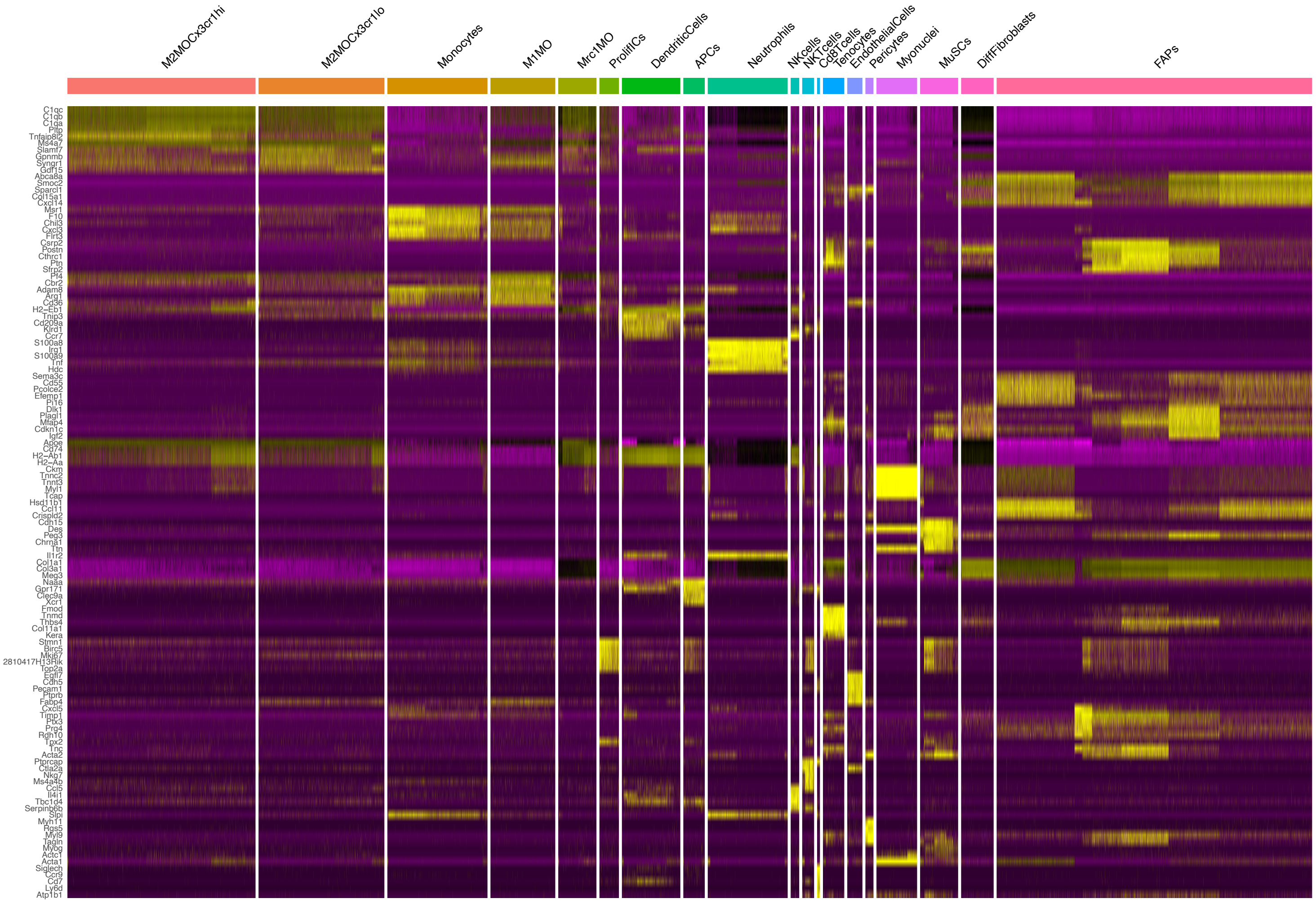


**Supplementary figure 4. Heat map plot of top 10 upregulated genes and corresponding annotated cell populations.** (A) Heat map plot showing distinct and major grouped cell populations across skeletal muscle homeostasis and regeneration. Cell populations and subpopulations were ordered based on the *top 10* expressed genes in each cluster subset shown in Supplementary figure 3. Please, note a high degree of transcriptomic variation among the FAPs subcluster.
