## Supplemental Figure 5 for "Single-Cell Transcriptome Dynamics of the Autotaxin-Lysophosphatidic Acid Axis During Muscle Regeneration Reveal Proliferative Effects in Mesenchymal Fibro-Adipogenic Progenitors"

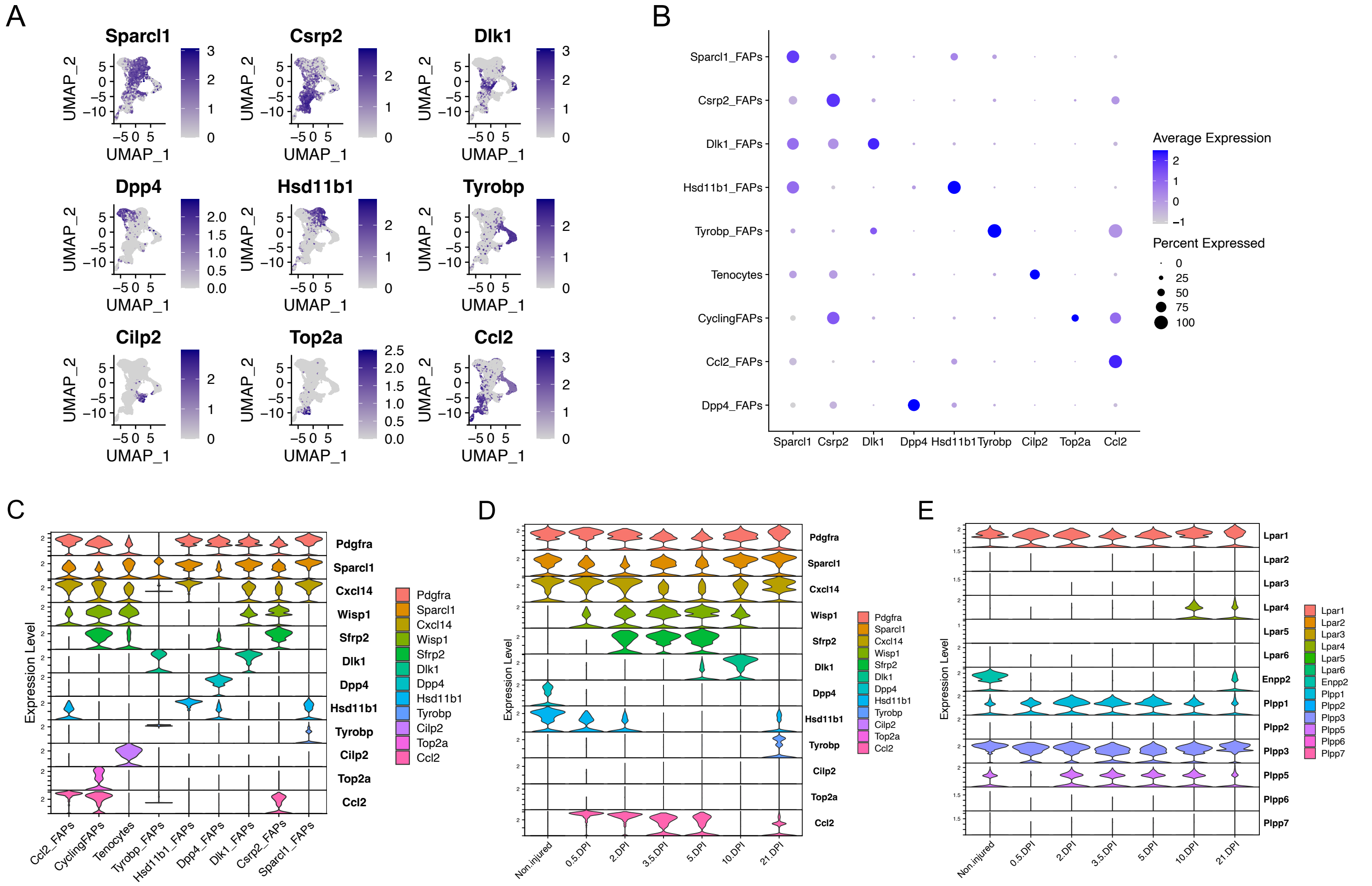


**Supplementary figure 5. Subclustering of the fibro-adipogenic progenitor and tenocyte lineages.** (A) UMAP plots showing 9 distinct and differentially expressed genes between FAPs and tenocyte (*Cilp2* expressing) populations across skeletal muscle homeostasis and regeneration when subclustering FAPs, DiffFibroblasts (i.e., Tyrobp_FAPs), and tenocytes subclusters. (B) Dot plot showing gene expression levels of the 9 differentially expressed genes shown in (A) between FAPs and tenocyte (*Cilp2* expressing) populations. (C) Violin plots showing the gene expression level of several FAP and tenocyte marker genes across the different FAPs and tenocyte subclusters. (D) Violin plots showing the gene expression level of several FAP and tenocyte marker genes across different conditions (uninjured and injured muscle) at different time points. DPI: days post-injury. (E) Violin plots showing the gene expression levels of LPAR, Enpp2, and Plpp family members across different conditions. Note that *Enpp2*/ATX gene is drastically downregulated upon injury.
