## Supplemental Figure 6 for "Single-Cell Transcriptome Dynamics of the Autotaxin-Lysophosphatidic Acid Axis During Muscle Regeneration Reveal Proliferative Effects in Mesenchymal Fibro-Adipogenic Progenitors"

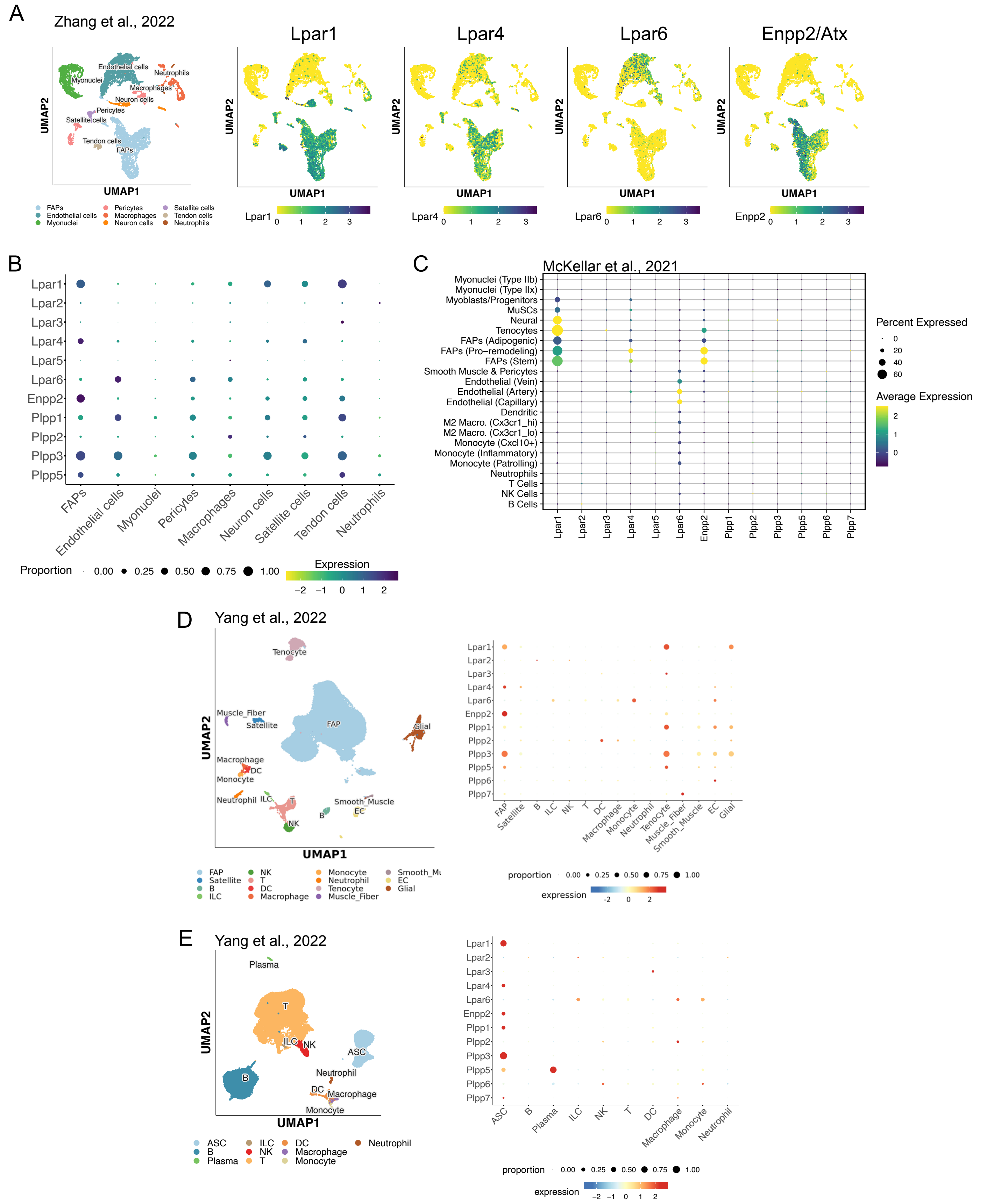


**Supplementary Figure 6. Validation of the ENPP2-LPAR-PLPP axis gene expression using different transcriptomic datasets.** (A) UMAP plot of scRNA-seq public data (Zhang et al., 2022) showing 9 distinct cell lineages in single cells across skeletal muscle homeostasis. Detected major cell lineages were colored by the predominant cell type(s) that composes each cluster. (B) Dot plots showing the expression level of several ATX-LPAR-PLPP genes across the different cell populations depicted in (A). (C) Dot plots showing the expression level of several ATX-LPAR-PLPP genes across the different cell populations as described in McKellar et al. 2021. (D) UMAP and dot plots showing the expression level of several ATX-LPAR-PLPP genes across the different populations as described in Yang et al., 2022. (E) UMAP and dot plots showing the expression level of several ATX-LPAR-PLPP genes across the different populations in adipose tissue as described in Yang et al., 2022.
