## Supplemental Figure 7 for "Single-Cell Transcriptome Dynamics of the Autotaxin-Lysophosphatidic Acid Axis During Muscle Regeneration Reveal Proliferative Effects in Mesenchymal Fibro-Adipogenic Progenitors"

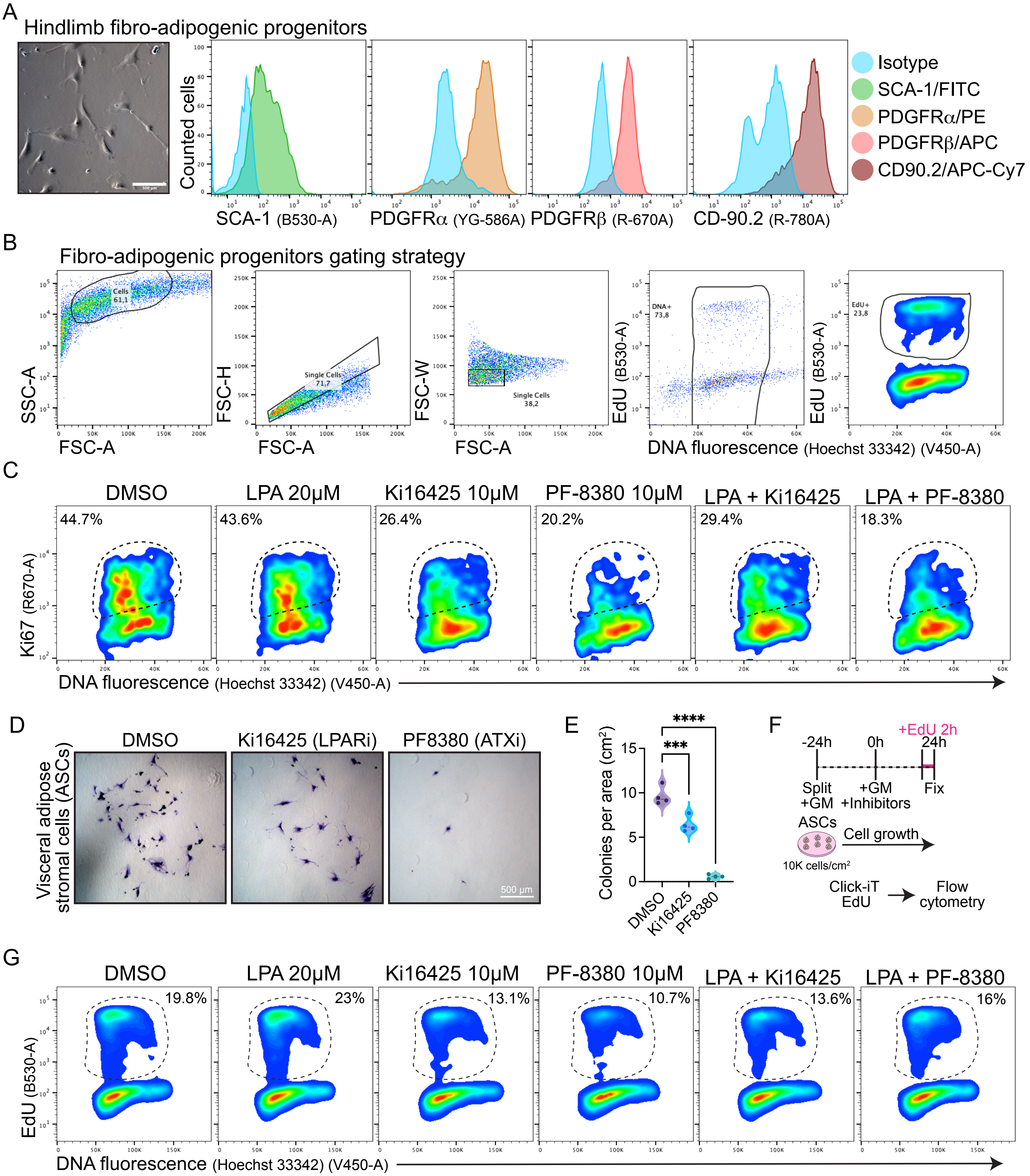


**Supplementary Figure 7. Pharmacological inhibiton of LPA receptors and Autotaxin reduces adipose stromal cell growth and proliferation.** (A) Brightfield image of cultured FAPs. Histograms showing the fluorescence intensity per counted FAPs at day 7 of cell growth (without passaging). (B) Representative flow cytometry plots showing the gating strategy used for EdU detection in muscle FAPs. (C) Flow cytometry detection of Ki67 labelled cells in combination with DNA fluorescence at 24 h of treatments. (D) Representative images of ASCs control-treated (DMSO) or treated with Ki16425 (10µM, LPAR1/3 antagonist) and PF-8380 (10µM, ATX inhibitor) as previously shown in Fig. 6A, and then stained with Crystal Violet. Scale bar: 500 µm. (E) Quantification of the number of cells per area as shown in (D) from four independent experiments.***P < 0.0002 ****P < 0.0001 by one-way ANOVA with Tukey's multiple comparison post-test; n = 4. (F) Outline of EdU assay using ASCs. (G) Flow cytometry detection of EdU labelled cells in combination with DNA fluorescence at 24 h of treatments.
