## Supplemental Figure 8 for "Single-Cell Transcriptome Dynamics of the Autotaxin-Lysophosphatidic Acid Axis During Muscle Regeneration Reveal Proliferative Effects in Mesenchymal Fibro-Adipogenic Progenitors"

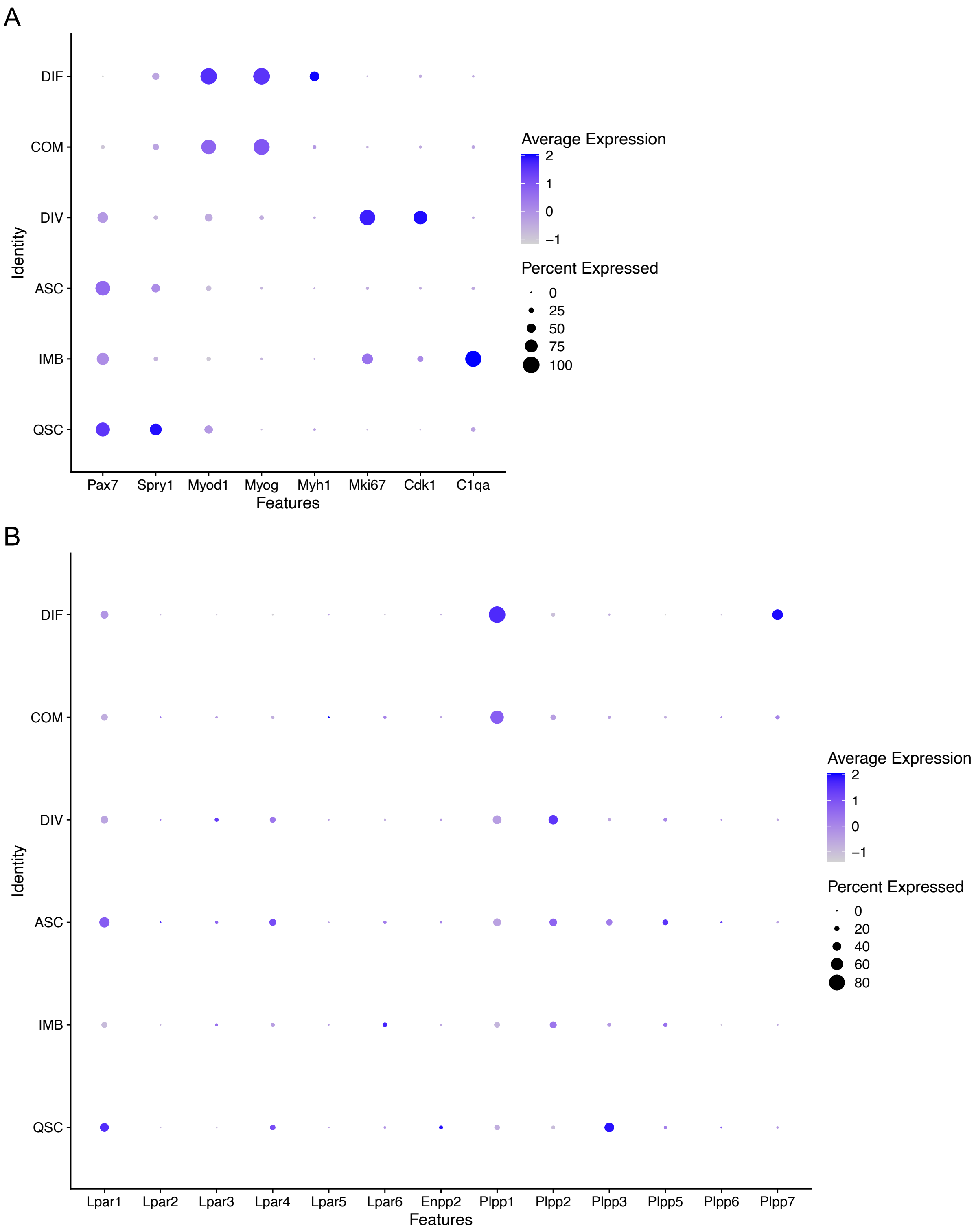


**Supplementary figure 8. Gene expression dynamics of the ATX-LPAR-PLPP axis in muscle stem cells.** (A) Dot plot showing gene expression levels of 8 differentially expressed genes between MuSC subpopulations. (B) Dot plot showing gene expression levels of LPAR, Enpp2, and Plpp family members in distinct MuSC subpopulations.
